# Genetic architecture of acute hyperthermia resistance in juvenile rainbow trout (Oncorhynchus mykiss) and genetic correlations with production traits

**DOI:** 10.1101/2022.11.14.516387

**Authors:** Henri Lagarde, Delphine Lallias, Pierre Patrice, Martin Prchal, Yoannah François, Jonathan D’Ambrosio, Emilien Segret, Ana Acin-Perez, Frederic Cachelou, Pierrick Haffray, Audrey Dehaullon, Mathilde Dupont-Nivet, Florence Phocas

## Abstract

**Background:** Selective breeding is a promising solution to reduce fish farms vulnerability to heat peaks which intensity and frequency are predicted to increase due to climate change. However, limited information about the genetic architecture of acute hyperthermia resistance in fish is available.

Two batches of sibs from a rainbow trout commercial line were produced. The first batch (N=1,382) was phenotyped for acute hyperthermia resistance at nine months, and the second batch (N=1,506) was phenotyped for main production traits (growth, body length, muscle fat content and carcass yield) at twenty months. Fish were genotyped on a 57K SNP array, and their genotypes were imputed at high-density thanks to their parents being genotyped on a 665K SNP array.

**Results:** The heritability estimate of resistance to acute hyperthermia in juveniles was 0.29 ± 0.05, confirming the potential of selective breeding for this trait. Genetic correlations between acute hyperthermia resistance and main production traits at near harvest age were all close to zero. Hence, selecting for acute hyperthermia resistance should not impact the main production traits, and reversely.

The genome-wide association study revealed that resistance to acute hyperthermia is highly polygenic; altogether, the six detected QTL explained less than 5% of the genetic variance. Two of these QTL, including the most significant one, might explain acute hyperthermia resistance differences across INRAE isogenic lines of rainbow trout. The phenotypic mean differences between homozygotes at peak SNP were up to 69% of the phenotypic standard deviation, showing promising potential for marker-assisted selection. We identified 89 candidate genes within the six QTL regions, among which the most convincing functional candidate genes were *dnajc7*, *hsp70b*, *nkiras2*, *cdk12*, *phb*, *fkbp10*, *ddx5*, *cygb1*, *enpp7*, *pdhx* and *acly*.

**Conclusions:** This study provides valuable insight on the genetic architecture of acute hyperthermia resistance in juvenile rainbow trout. The potential for the selective breeding of this trait was shown to be substantial and should not interfere with selection for main production traits. Identified functional candidate genes give a new insight on physiological mechanisms involved in acute hyperthermia resistance, such as protein chaperoning, oxidative stress response, homeostasis maintenance and cell survival.

## Background

Aquaculture, which produced 7.5% of animal-source proteins consumed worldwide in 2018, is a significant contributor to world food security [1]. The importance of aquaculture in global food production is expected to strengthen in the medium term as it grows faster than the other main food production sectors, with a mean annual growth of 4.6% between 2010 and 2020 [2]. However, this steady increase in aquaculture production over the past few decades could be jeopardised in the future by the effects of climate change. These effects are expected to be multiple such as the chronic water temperature increase, more frequent and severe extreme weather events or changes in rainfall patterns affecting all levels of production by reduced growth and survival, new and more frequent epidemics or input shortages in fish farms [3].

As ectotherms, fish are particularly susceptible to changes in water temperature. Hence, the effects of climate change, both through chronic temperature increases and acute hyperthermia conditions induced by more frequent and intense heat waves, are expected to substantially impact fish production [4]. Extreme temperature events are predicted to impact ectotherms populations more than climate warming [5]. Furthermore, the adaptative capacities of the European perch, *Perca fluviatilis*, were found to be higher for resistance to chronic hyperthermia stress compared to resistance to acute hyperthermia stress [6].

Hence, improving fish farms resilience to heat wave events is a primary concern. Acute hyperthermia stress causes a series of physiological and behavioural changes in fish, starting with catecholamines and cortisol releases, followed by an increase in plasma glucose and lactate, the over-expression of genes associated with acute hyperthermia stress such as heat shock proteins and the lowering or cessation of feeding [4]. The consequences of acute hyperthermia on production efficiency in fish farms are significant, among which growth losses and high mortalities (e.g. [7]). Adaptation strategies must therefore be established to guarantee food security by limiting production losses induced by heat waves. Promising strategies include engineering solutions, species diversification, better farm management, nutrition, exercise training or genetic improvement [8, 9]. Genetic improvement by selective breeding is an interesting solution to enhance fish resistance to acute hyperthermia conditions as genetic gain is cumulative and can be quickly disseminated in farms thanks to the high fecundity of fish.

Rainbow trout *Oncorhynchus mykiss* is the second most farmed salmonid species with 959,600 tons yielded in the world in 2020 [2]. Like other salmonids, rainbow trout is particularly sensitive to acute hyperthermia stress as its upper thermal resistance, the temperature at which it loses equilibrium, is usually between 26°C and 30°C [10, 11]. Breeding rainbow trout robust to acute hyperthermia stress would help fish farmers adapt to climate change.

The heritability, the genetic correlations with production traits and the identification of quantitative traits loci (QTL) are needed to optimise and assess the profitability of genetic improvement by selective breeding of a new trait.

Heritability is the key information for estimating the expected genetic gain at each generation and thus to assess the theoretical cost-benefit of selecting the given trait [12]. Heritability of acute hyperthermia resistance was estimated to be 0.41 ± 0.07 in a North-American population of rainbow trout, demonstrating an interesting potential for genetic improvement of this trait [13]. Nevertheless, the realised heritability was only 0.10 ± 0.05 in six generations of selection for acute hyperthermia resistance in the zebrafish *Danio rerio* [14]. Low heritability could question the genetic lever’s relevance to improve fish resistance to acute hyperthermia stress.

Estimating genetic correlations is also essential to evaluate the effect of selection for one trait on other traits of interest and vice-versa [12]. Three of the main traits of interest in rainbow trout breeding programs are growth, carcass yield and fillet fat percentage at harvest age [15]. Estimating genetic correlations between these traits and acute hyperthermia resistance is all the more critical as trade-offs were suggested between growth and acute hyperthermia resistance in rainbow trout [11, 16]. The genetic correlation between acute hyperthermia resistance and growth was estimated to be -0.03 ± 0.18 in a North-American commercial population of rainbow trout [13]. However, no similar study was realised on European commercial populations. Moreover, in Perry et al. [13], fish were less than one year old, which is still far from the harvest age required for rainbow trout filleting. Eventually, to our knowledge, genetic correlations between acute hyperthermia resistance and carcass yield or fillet fat percentage were never estimated in any fish species at any age.

Detecting QTL enables the identification of genetic markers tightly associated with the phenotype of interest. These genetic markers can then be used to improve the genomic prediction through weighted GBLUP [17–20] or for marker-assisted selection as it was successfully performed for improving resistance to infectious pancreatic necrosis disease in Atlantic salmon *Salmo salar* [21–23]. Moreover, searching for candidate genes in QTL regions allows to better understand the underlying mechanisms involved in acute hyperthermia resistance. Pioneering works studied the genetic architecture of acute hyperthermia resistance in the same North-American population of rainbow trout as in Perry et al. [13] and found QTL but with large confidence intervals due to the low density of markers [24–27]. More recent studies using high-density genotyping chip showed that acute hyperthermia resistance is a polygenic trait and identified genes associated with this trait in channel catfish *Ictalurus punctatus* and in large yellow croaker *Larimichthys crocea* [28, 29]. Such study has not yet been conducted on rainbow trout.

The objective of the present study was to present a complete overview of the genetic architecture of acute hyperthermia resistance in a French commercial line of rainbow trout, never studied for this trait, taking advantage of the recent advances in genomics. For this purpose, a group of fish issued from 76 females mated with 99 males in ten independent full-factorial blocks families was phenotyped at nine months for acute hyperthermia resistance and their sibs were phenotyped at twenty months, which is near harvest age (body weight around 1 kg), for production traits using a robust experimental design with over 1,300 phenotyped fish in each group. Phenotyped fish were genotyped for 57K SNP (Single Nucleotide Polymorphism markers) [30] and imputed to 665K SNP thanks to their parents being genotyped with a new high-density SNP array [31]. The main issues addressed in this study were to i) assess genetic parameters for acute hyperthermia resistance in juveniles, ii) estimate genetic correlations between acute hyperthermia resistance and production traits at harvest age, iii) detect with a powerful tool (665K SNP array) QTL associated with acute hyperthermia resistance and (iv) identify functional candidate genes.

## Methods

### Fish production

The fish production process is summarised in Figure 1. Fish were derived from a commercial line from ’Viviers de Sarrance’ (Sarrance, France) breeding company which had been mass selected for growth, morphology according to a salmon-like shape and gutted carcass yield with ultrasound during nine generations [32] and, more recently, for the body colour and carcass yield based on sib selection. In November 2019, 76 two-year-old females were mated with 99 neomales (sex-reversed XX females used as sires) in ten independent full-factorial blocks of 9- 10 sires crossed with 7-8 dams in the ’Labedan’ hatchery (64490 Sarrance, France). Fin samples were collected for later genotyping of all parents. Fertilised eggs were all mixed and incubated in a 40-litres cylindrical-conical incubator at 8.5-9°C. At 29 days post-fertilisation (dpf), eyed-stage eggs were transferred to ’Les fontaines d’Escot’ fish farm (64490 Sarrance, France). Eggs were randomly placed in baskets in flow through hatching troughs containing 13-14°C circulating water for hatching. Hatching occurred between 29 dpf and 36 dpf. At 101 dpf, about 4,500 fries were randomly transferred in two nursery tanks supplied with the same water source. At 127 dpf, fish were transferred to the ’Viviers de Rébénacq’ fish farm (64260 Rébénacq, France), located forty kilometres from the previous farm. Fish from the two nursery tanks were grouped and reared in a 23m^3^ concrete raceway. At 265-dpf, fish were randomly divided into two batches of approximately equal size named batch 1 (B1) and batch 2 (B2). Fish from B1 were tagged (Biolog-id, 1.4 × 8 mm) and fin-clipped for later DNA analysis. B1 fish were then placed in a second concrete raceway for a week before being transferred to ANSES-SYSAAF Fortior Genetics platform (Brittany, France) for acute hyperthermia resistance phenotyping. Batch 2 (B2) was made of all remaining fish and B2 fish stayed in the 23m^3^ concrete raceway.

**Figure 1:**
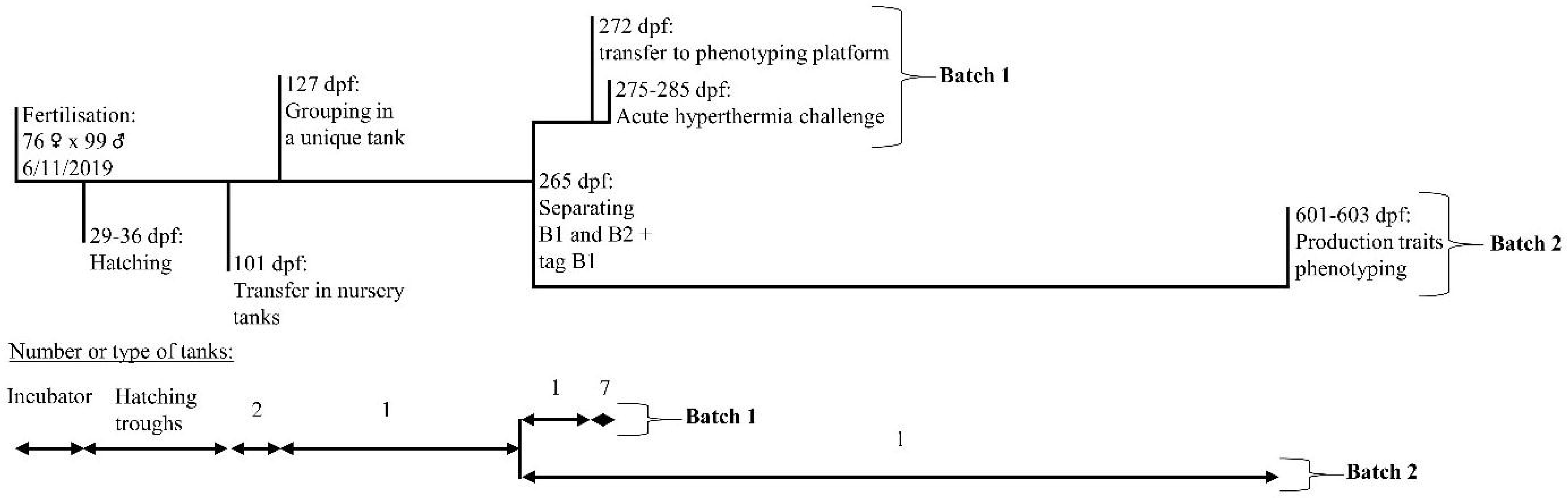
**Batch 1 (B1, juveniles) and Batch 2 (B2, harvest size) rainbow trout production processes. Abbreviations: dpf, day post-fertilisation.**

At 425 dpf, B2 fish (mean body weight of 300g) were transferred in a 93m^3^ concrete raceway and stayed in until phenotyping for production traits (between 600 dpf and 604 dpf).

During growth, fish were fed *ad libitum* using extruded commercial feed (Le Gouessant, Lamballe, France). At ’Viviers de Rébénacq’ fish farm, fish were exposed to a minimum water temperature of 9.8°C in February 2020 and a maximum of 14.5°C in August 2020. Minimum oxygen concentration measured was 8.7mg/L in August 2020. Density was between 6kg/m^3^ and 24kg/m^3^ in both batches. Survival rate from hatching to transfer of B1 to ANSES-SYSAAF Fortior Genetics platform was 65% and survival rate from hatching to phenotyping of B2 was 63%.

### Phenotyping

#### Phenotyping for acute hyperthermia resistance (batch B1)

At 272 dpf, B1 fish (N=1,382) were transferred to ANSES-SYSAAF Fortior Genetics platform (Plouzané, France) for acute hyperthermia resistance phenotyping. Transportation was done by truck in controlled conditions (10h of transportation, density: 40kg/m^3^, temperature: 12 ± 1°C, O2 concentration around 12.9mg/L). Once arrived at the platform, fish were randomly distributed in seven fibreglass 350L tanks with 197 ± 12 fish per tank for acclimation. Tanks were supplied with filtrated and sterilised river water with a daily temperature range of 16°C- 18°C. Oxygen saturation level was maintained high by continuously and softly bubbling compressed air. The phenotyping was done by group, the seven fibreglass tanks corresponding to the seven phenotyping groups. It was performed at the rate of one group per day, spread between 275 dpf and 285 dpf. Accordingly, the duration of acclimation, i.e. the time between the arrival at the phenotyping platform and the phenotyping day, was between two to twelve days depending on group phenotyping order. The duration of acclimation will be discussed further. Each group was starved 24 hours before the phenotyping challenge.

Acute hyperthermia challenge was conducted in the following way: at 9 a.m., fish from a group were transferred with a landing net to the challenge tank (350L, fibreglass), supplied with the same river water as in the acclimation tanks. Once the transfer completed, temperature was gradually increased from the initial challenge tank water temperature (17.3 ± 0.7 °C, min: 16.1°C in group 7 (G7), max: 18.4°C in G4) at a rate of 3.1 °C/hour during the first 1.5 hours and then at a rate of 0.9 °C/hour during the rest of the challenge by adding heated water from a buffer tank. The heating curves experienced by the seven groups are presented in Figure 2. Oxygen saturation was maintained above 80% by softly bubbling pure O2. Above 27°C, fish gradually began to lose equilibrium. Each fish losing equilibrium was removed from the tank and identified by reading its tag. The time was recorded and the fish was weighed and euthanised by overdose of anaesthetic (Eugenol, 180 mg/L). Challenges ended when the last fish lost its equilibrium. Temperature, O2 concentration and O2 saturation were recorded every ten minutes using electronic probes (OxyGuard, Handy Polaris). NH4+ concentration, pH and CO2 concentration were checked in the first three groups at the beginning and in the middle of the challenge with kits for NH4+ (Tetra, Test), NH3/NH4+ and pH (JBL, pH test) and a CO2 analyser (Oxyguard, CO2 Portable).

**Figure 2:**
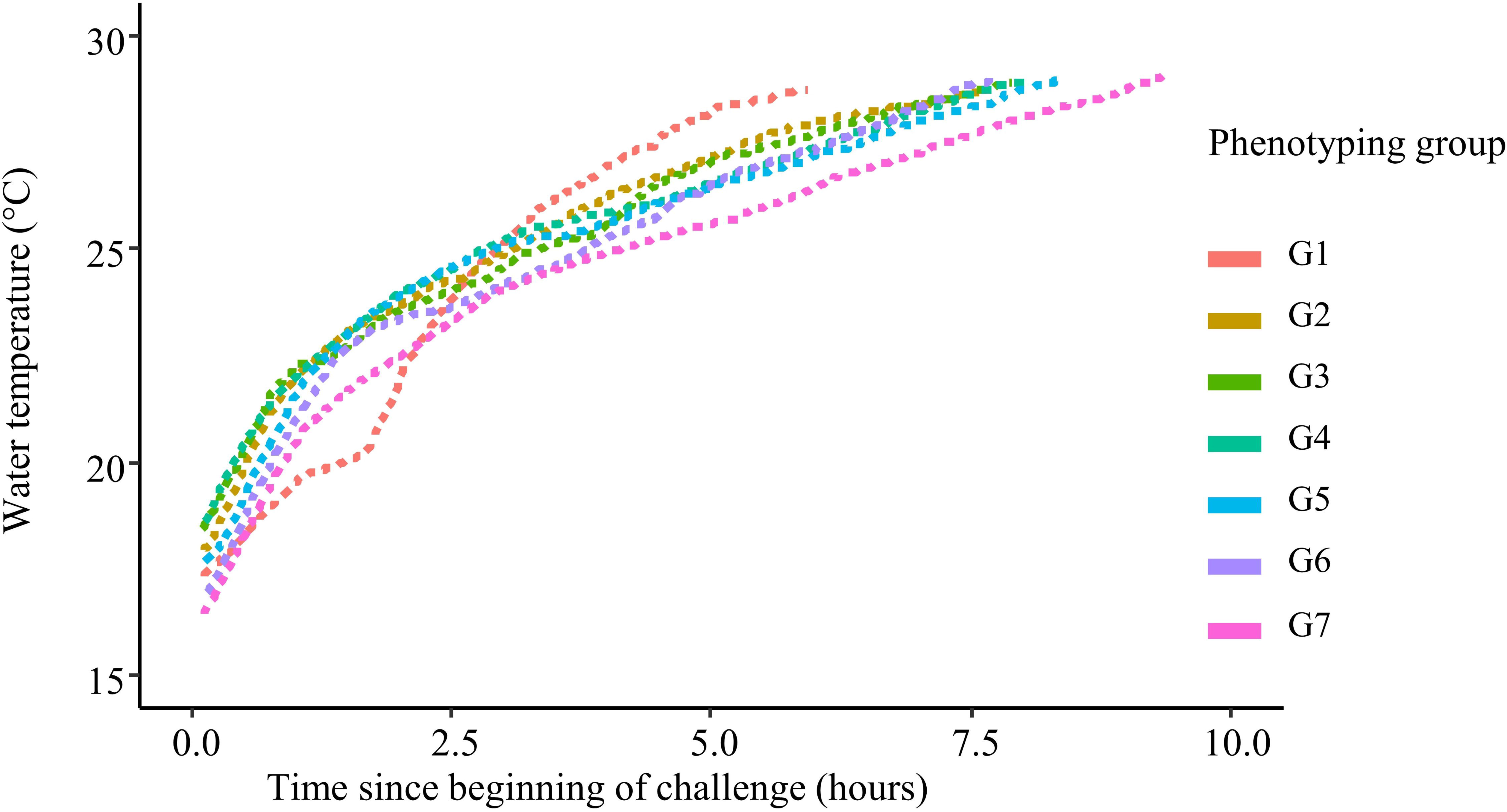
**Kinetics of temperature increase in the seven rainbow trout groups of batch 1 (B1, juveniles) batch of acute hyperthermia challenge.**

From B1 batch, we recorded acute hyperthermia resistance and the body weight named BW1. Raw acute hyperthermia resistance phenotype was measured as the raw time to loss of equilibrium (rTLE). For reasons presented in the results section, rTLE was centred to 0 and reduced to 1 within each phenotyping day. This standardised phenotype was named TLE.

#### Phenotyping for production traits at commercial size (batch B2)

Fish from B2 batch (N=1,506) were phenotyped for production traits at commercial size in three days between 601 dpf and 603 dpf. Every day, 502 ± 49 fish were slaughtered by bleeding in icy water immediately after netting. Fish were slaughtered by group of 20 fish and stored on ice to avoid rigor mortis adverse effects.

Once killed, fish were processed and phenotyped in thirty minutes maximum. A fin sample was first collected on each fish for later genotyping and kept in 90 % ethanol. The following raw phenotypes were then measured: body weight (BW2), fork length (FL), head weight (HeadW), headless gutted carcass weight (HGCW) as an indirect measurement of the fillet yield [33] and viscera weight (ViscW). Fillet fat percentage (Fat%) was estimated by micro-waves using a Distell Fish-Fatmeter®. The Fatmeter was applied at the anterior and posterior dorsal positions above the lateral line of the fish’s left side [34]. Fat% was the mean of these two measurements.

All weights were measured to the nearest 0.5 g, fork length to the nearest 0.5 mm and total fat to the nearest 0.1%. When difference between BW2 and the sum of HGCW, HeadW and ViscW was more than 10 grams, fish BW2 and HGCW data were removed from the dataset (N=24 individuals).

We analysed body weight (BW2), fork length (FL), fillet fat percentage (Fat%) and headless gutted carcass yield (HGC%) calculated as the ratio of HGCW to BW2.

### Genotyping

The collected fin samples from the 99 sires, the 76 dams, 1,382 offspring of B1 and 1,506 offspring of B2, were genotyped by INRAE genotyping platform Gentyane (Clermont-Ferrand, France). The 2,888 offspring of B1 and B2 were genotyped for 57,501 SNP (medium-density (MD) genotypes) with the 57K SNP AxiomTM Trout Genotyping array from Thermo Fisher [30]. The 175 parents were genotyped for 664,531 SNP (high density (HD) genotypes) with the 665K SNP AxiomTM Trout Genotyping array from Thermo Fisher [31].

Among the genotyped individuals, eight offspring from B2 and one sire had less than 90% of the SNP genotyped and their genotypes were therefore removed from the analysis. SNP with probe polymorphism and multiple locations on the Arlee genome assembly (accession number: GCA_013265735.3; [35]) were also discarded from the analysis as described by Bernard et al. [31]. In addition, only SNP with a call rate higher than 0.97, a test of deviation from Hardy-Weinberg equilibrium with a p-value > 0.00001 and a minor allele frequency higher than 0.05 were kept for the study. The number of SNP passing quality control filters was 30,325 in MD genotypes and 420,079 in HD genotypes.

Parentage assignment was done using 1,000 SNP from the MD chip equidistantly distributed across the genome with the R package APIS [36] with a positive assignment error rate set to 5%. A total of 55 fish were unassigned in B1 and 48 in B2, probably due partly to the non-genotyped sire. In B1, the mean number of phenotyped and genotyped progenies per sire, per dam and full-sibs family were respectively 13.5 ± 5.1, 18.4 ± 6.5 and 2.4 ± 1.4 (unassigned individuals excluded). In B2, the mean number of phenotyped and genotyped progenies per sire, per dam and full-sibs family were respectively 14.8 ± 5.3, 20.4 ± 8.5 and 2.5 ± 1.5 (unassigned individuals excluded).

As explained in the data analysis part, maternal effect was significant for acute hyperthermia resistance phenotype. Unassigned individuals were therefore removed from the dataset, making a total of 1,327 analysed fish in B1. Conversely, maternal effect was not significant for phenotypes measured in B2, unassigned individuals were therefore kept. The total number of analysed individuals in B2 was 1,498 (1,506 minus the eight badly genotyped individuals).

Utilising quality-filtered genotypes and pedigree information, FImpute3 software [37] was used to impute missing genotypes in both offspring and parents and to impute offspring’s MD genotypes to HD ones thanks to the parental reference HD genotypes.

### Data analysis

#### Estimation of genetic parameters

Genetic (co)variance components were estimated for all traits with AIREML algorithm in BLUPF90 software [38] using the following animal model:

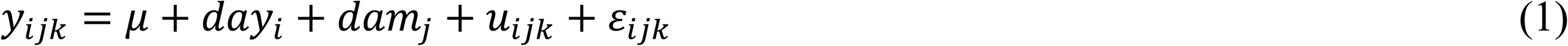

where *y_ijk_* is the performance of animal k, μ is the overall mean of the population, *day_i_* is the fixed effect of day of challenge i (day/group of phenotyping for batch 1, day of slaughtering for batch 2), *dam_j_* is the random effect of dam j, *u_ijk_* is the additive genetic effect of animal k and *ε_ijk_* is the random residual error. Both pedigree and genomic relationship matrices were computed. The pedigree was constituted of 20,372 animals over ten generations. The genomic matrix was built with imputed HD genotypes [39].

Heritability of each trait was calculated using univariate analysis with both pedigree and genomic relationship matrices as:

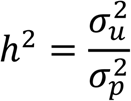

where 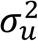 is the additive genetic variance and 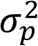 is the phenotypic variance.

The genetic correlation (*r_g_*) between two traits x and y was calculated using bivariate analyses with both pedigree and genomic relationship matrices:

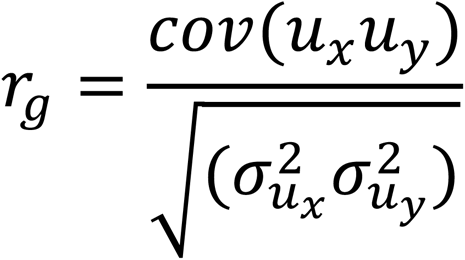

where *cov*(*u_x_u_y_*) is the additive genetic covariance between x and y, and 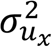 and 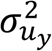 are the additive genetic variance of x and y.

#### Genome-Wide Association Study and QTL Detection for acute hyperthermia resistance

A Bayesian variable selection model with a Bayes Cπ approach was used to locate QTL and estimate the proportions of genetic variance explained by the identified QTL [40]. The marker effects were estimated through the Markov Chain Monte Carlo (MCMC) algorithm. At each cycle of the MCMC algorithm, only a given fraction (π) of the 420K markers were assumed to have a non-zero effect on the phenotype and to follow a normal distribution 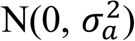, with 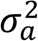 the additive genetic variance. The remaining fraction (1-π) of markers have zero-effect.

The model was the following:

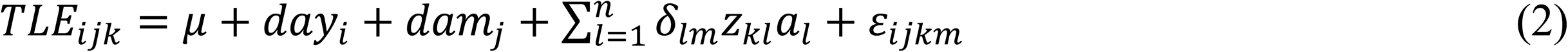

With *TLE_ijk_*, the acute hyperthermia resistance phenotype of animal k, *µ* the overall mean of the population, *day_i_* the fixed effect of day of challenge i, *dam_j_* the random effect of dam j and n the total number of SNP (420,079). *δ_lm_* is an indicator variable: within a cycle m, *δ_lm_*=1 if the effect of SNP l is estimated in this cycle or *δ_lm_*=0 otherwise. In each cycle, *δ_lm_* is sampled from a binomial distribution with a probability π that *δ_lm_* was equal to 1. The proportion π was sampled from a beta distribution B(α,β) with α = 400 and β =420,079; it approximatively corresponds to 400 SNP selected at each cycle with a non-zero effect. *δ_lm_* was the genotype on locus l for individual k (coded as 0, 1, or 2), *a_l_* the effect of the reference allele of SNP l, and *ε_ijkm_* the residual effect. The Markov chain Monte Carlo was run with 600,000 cycles and a burn-in period of 10,000 cycles. Results were saved every 40 cycles.

Convergence was assessed by two approaches. First, plots of the posterior density of genetic and residual variances were verified by visual inspection. Secondly, genomic breeding values estimated from two differently seeded runs of the MCMC algorithm were highly correlated (r > 0.99).

The Bayes factor (BF) was calculated to quantify the degree of association between a SNP and the resistance to acute hyperthermia. For the i^th^ SNP, the BF is equal to Pi/(1−Pi) π/(1−π), with Pi the probability of the SNP to be included in the model as having a non-zero effect and π, the given fraction of the 420K markers assumed to have a non-zero effect on the phenotype. As proposed by Kass et al. [41], a logBF value (derived as twice the natural logarithm of BF) greater than or equal to a threshold of 6 at a peak SNP was used as strong evidence for the existence of a QTL. Following Michenet et al. [42], a credibility interval was computed around the peak SNP of each QTL including all SNP for which logBF≥3 in a sliding window of 200 kb on both sides of the peak SNP. Genes within QTL credibility intervals were annotated with the NCBI *O. mykiss* Arlee genome assembly (GCA_013265735.3 USDA_OmykA_1.1; [35]).

#### TLE depending on genotypes at SNP peaks

We analysed individuals’ acute hyperthermia resistance phenotypes depending on their genotypes at the peak SNP of detected QTL. Significance of the difference in TLE between the two homozygous genotypes at each peak SNP was analysed by Anova Tukey tests. The difference was expressed in percentages of the phenotypic standard deviations. Dominance effect was quantified at each SNP peak as the difference between TLE of the heterozygote genotype and the average of the two homozygous genotypes mean TLE. Significance of dominance effect was tested using one sample t-test. Statistical tests were considered significant at an alpha= 0.05.

#### Pre-validation of identified QTL in isogenic lines of rainbow trout

In a previous experiment, we measured the acute hyperthermia resistance phenotypes of six isogenic lines of rainbow trout at 185 dpf (6 months) and 457 dpf (15 months) using a protocol similar to the one used in the present study [16]. Isogenic lines are powerful experimental genetic resources. Within isogenic lines, fish share the same genotype while the different lines represent a sample of the genetic variability of the INRAE synthetic population from which they were derived [43].

At 185 and 457 dpf, the isogenic line named A32h was found to be the most resistant and the isogenic line named A22h was found to be the most sensitive [16]. Consequently, A32h was called the resistant isogenic line and A22h the sensitive isogenic line for the current paper. The four other isogenic lines phenotyped in Lagarde et al. [16] either had an intermediate ranking or changed their position in the resistance ranking between 185 and 457 dpf and were therefore not considered here.

Isogenic lines A32h and A22h are heterozygous lines as they were produced by mating females from a unique homozygous isogenic line named B57 with males from homozygous isogenic lines A32 and A22. Therefore, heterozygous isogenic lines A32h and A22h shared the same maternal genetic basis but had different paternal genetic basis. Homozygous isogenic lines A32 and A22 have previously been sequenced to establish a catalogue of variants (Bernard et al., 2022). We only looked at the paternal genetic sequences A22 and A32 as maternal genetic sequence B57 was identical between A22h and A32h. Using these data already produced, we checked if some of the commercial population’s genetic polymorphisms associated with acute hyperthermia resistance were shared with isogenic lines. To do so, we searched for genetic polymorphism between A22 and A32 in the SNP previously identified to be strongly associated with TLE in the commercial population, i.e. which had logBF >6. For SNP meeting this criterion, we checked whether the resistant isogenic line held the favourable alleles (reference allele if *a_l_*>0 or the other allele if *a_l_*<0 for the SNP l) as estimated by GWAS in the commercial population.

## RESULTS

### Descriptive statistics of collected phenotypes

In the present study, two batches of all-female rainbow trout issued from the same families were phenotyped at 275-285 dpf for acute hyperthermia resistance (B1) or 600-604 dpf for production traits (B2). Descriptive statistics of collected phenotypes are presented in Table 1.

**Table 1.**
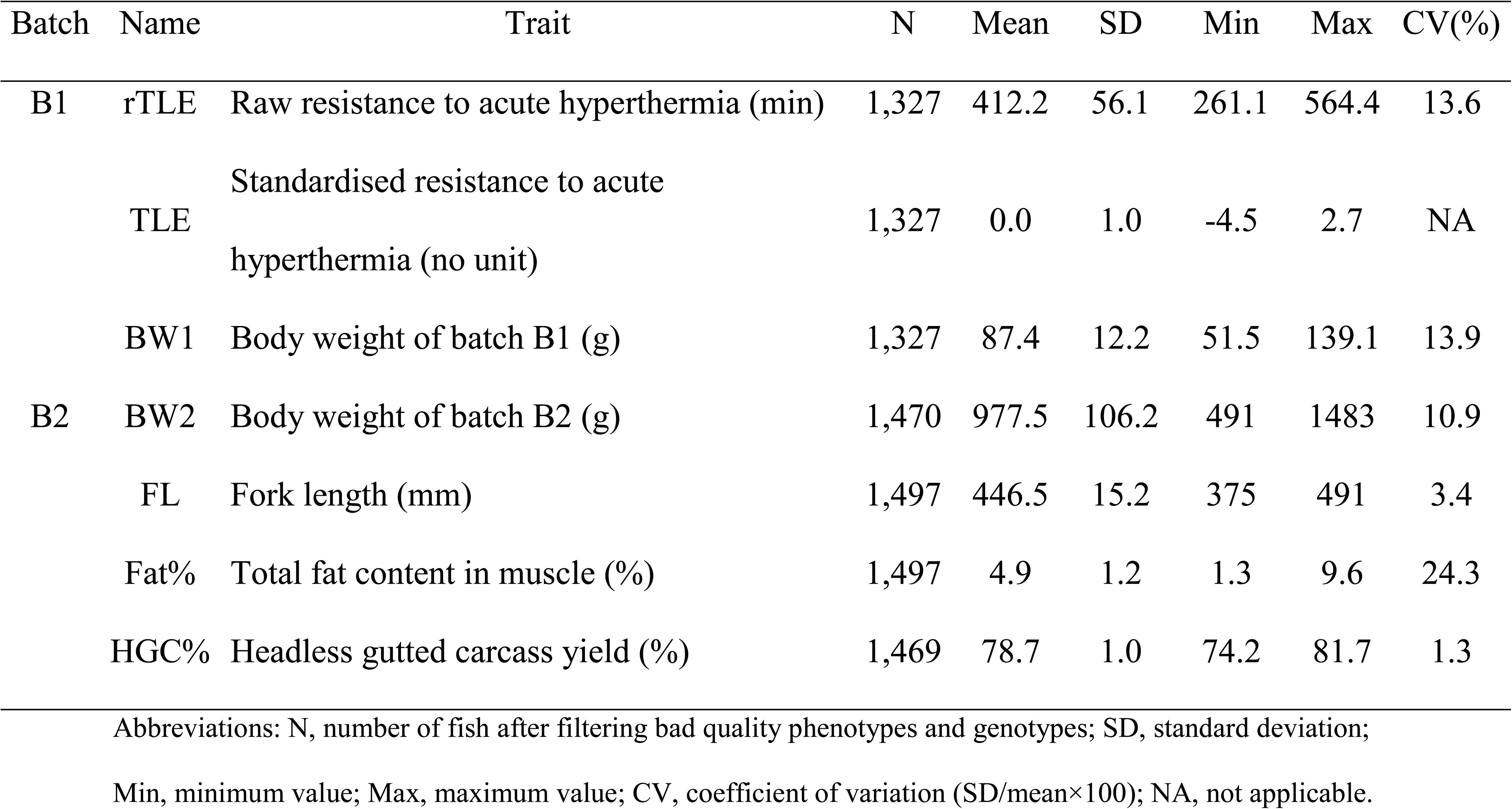
Descriptive statistics for acute hyperthermia resistance, growth, carcass yield and fat in rainbow trout.

| Batch | Name | Trait | N | Mean | SD | Min | Max | CV(%) |
| --- | --- | --- | --- | --- | --- | --- | --- | --- |
| B1 | rTLE | Raw resistance to acute hyperthermia (min) | 1,327 | 412.2 | 56.1 | 261.1 | 564.4 | 13.6 |
|  | TLE | Standardised resistance to acute hyperthermia (no unit) | 1,327 | 0.0 | 1.0 | -4.5 | 2.7 | NA |
|  | BW1 | Body weight of batch B1 (g) | 1,327 | 87.4 | 12.2 | 51.5 | 139.1 | 13.9 |
| B2 | BW2 | Body weight of batch B2 (g) | 1,470 | 977.5 | 106.2 | 491 | 1483 | 10.9 |
|  | FL | Fork length (mm) | 1,497 | 446.5 | 15.2 | 375 | 491 | 3.4 |
|  | Fat% | Total fat content in muscle (%) | 1,497 | 4.9 | 1.2 | 1.3 | 9.6 | 24.3 |
|  | HGC% | Headless gutted carcass yield (%) | 1,469 | 78.7 | 1.0 | 74.2 | 81.7 | 1.3 |

Abbreviations: N, number of fish after filtering bad quality phenotypes and genotypes; SD, standard deviation; Min, minimum value; Max, maximum value; CV, coefficient of variation ( $SD/mean \times 100$ ); NA, not applicable.

Mean BW1 was 85.6 ± 12.0 g in B1. Raw acute hyperthermia resistance was measured as the raw time to loss of equilibrium (rTLE). Mean rTLE was 412 ± 56 minutes, all phenotyping groups together. However, there were significant between-groups differences for rTLE mean (ANOVA, N=1,327, df=6; F value=402.7; P<2.2e-16) and rTLE’s standard deviation (Levene’s statistic, N=1,327, df=6; F value=56.9; P<0.001). As shown in Table 2, fish of G1 lost equilibrium very quickly and with low variability (mean of 319 ± 14 minutes), fish of G2, G3, G4, G5 and G6 showed intermediate and similar mean and SD (mean between 395 and 447 minutes, SD between 23 and 34 minutes) and fish of G7 lost equilibrium lately and with high variability (mean of 474 ± 62 minutes). The kinetics of loss of equilibrium is presented in Figure 3. To correct for these between-groups differences, rTLE was centred to 0 and reduced to 1 within each phenotyping day to obtain the corrected phenotype, TLE. Minimum and maximum TLE values were -4.5 and 2.7, respectively (Table 1).

**Figure 3:**
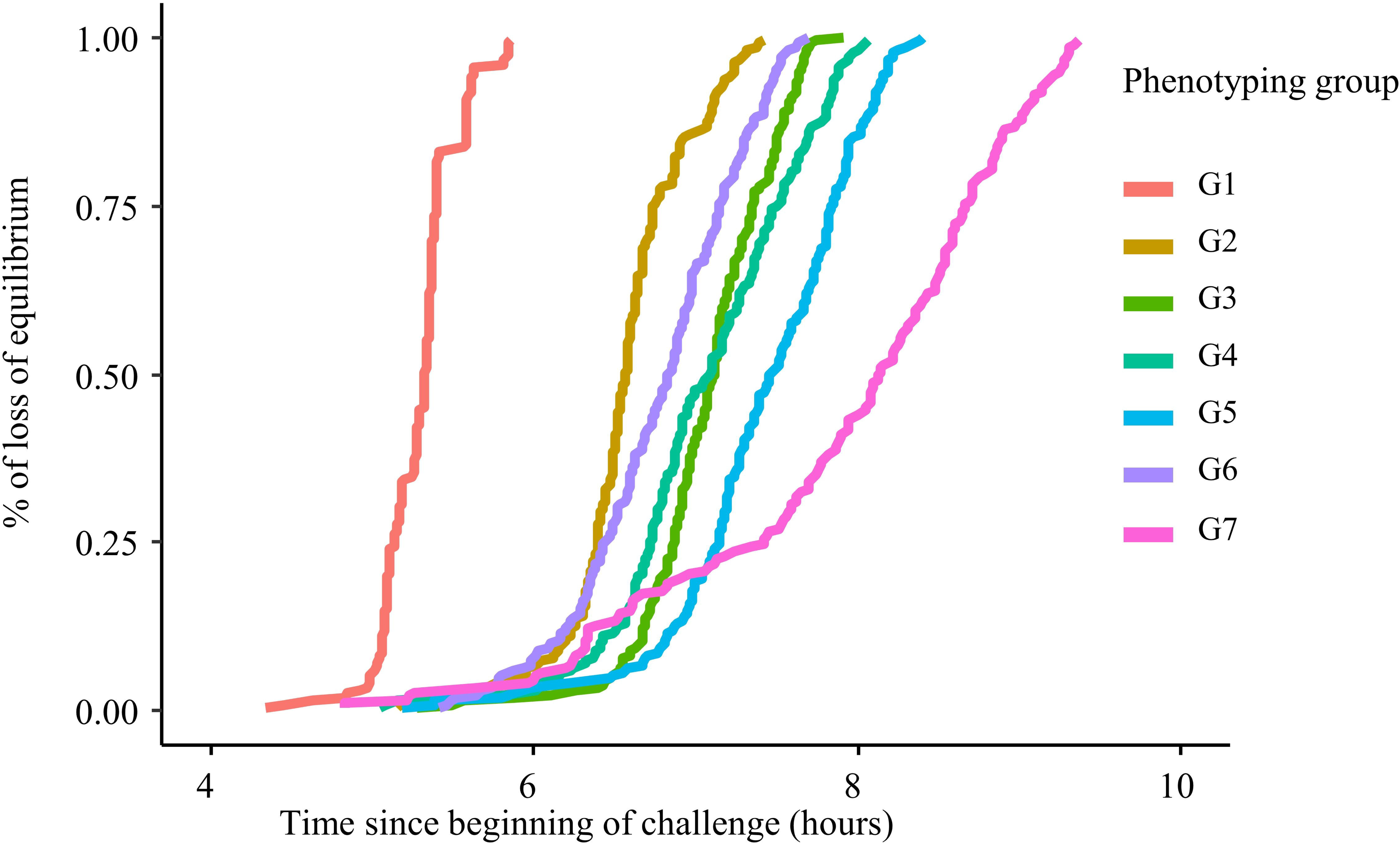
**Kinetics of cumulative loss of equilibrium in the seven rainbow trout groups of Batch 1 (B1, juveniles) of acute hyperthermia challenge.**

**Table 2.** Mean and standard deviation of raw TLE (rTLE) in acute hyperthermia phenotyping groups in B1.

| Phenotyping group | rTLE mean | rTLE SD | N |
| --- | --- | --- | --- |
| G1 | 318.7 | 14.1 | 187 (194) |
| G2 | 394.9 | 23.2 | 196 (202) |
| G3 | 424.8 | 24.4 | 192 (204) |
| G4 | 424.9 | 34.2 | 195 (200) |
| G5 | 446.8 | 34.0 | 199 (207) |
| G6 | 407.4 | 30.3 | 196 (202) |
| G7 | 473.8 | 61.6 | 162 (173) |

Abbreviations: rTLE mean, mean of raw resistance to acute hyperthermia (in minutes); rTLE SD, standard deviation of raw resistance to acute hyperthermia; N, number of fish phenotyped, genotyped and assigned. In brackets, number of fish phenotyped.

The number of acclimation days, defined as the days between the arrival of fish on the phenotyping platform and the phenotyping day of each group, and the temperature in the challenge tank at six hours were strongly correlated with these between-groups differences in mean and SD of rTLE. These two factors were confounded with a coefficient of correlation of -0.93: phenotyping groups with the highest acclimation days were also the ones with the lowest temperature in the challenge tank at six hours.

The number of days of acclimation varied from two days for G1 to twelve days for G7. The correlations between the number of acclimation day of each phenotyping group and the mean and SD of rTLE within these groups were 0.97 and 0.89, respectively. The temperature in the challenge tank six hours after the start of the challenge was also strongly correlated with TLE mean and SD with correlation coefficients of -0.94 and -0.94, respectively.

NH4+ concentration and pH were measured at precise time points in the three first groups. NH4+ concentration was under the threshold of colourimetric detection at the start of each challenge with a pH of 7.4 while between 0.25 and 1.5 mg/L in the three groups with a pH of 7.2±0.3 three hours after the start of the challenges, and between 2 and 3 mg/L in the three groups with a pH of 7.2±0.4 six hours after the start of the challenges. CO2 concentration was measured in the first two groups at the start of the challenge (9mg/L) and three hours after (12mg/L).

Descriptive statistics of phenotypes for processing traits at commercial size (B2) are given in Table 1. Briefly, mean and coefficient of variation were 977.5 g and 10.9% for BW2, 446.5 mm and 3.4% for FL, 4.9% and 24.3% for Fat% and 78.7% and 1.3% for HGC%.

### Genetic parameters for TLE and production traits

Heritability estimates and genetic correlations estimated with pedigree-based (Additional Table 1) or genomic information (Table 3) gave very similar results for all traits. GBLUP estimates were more accurate with lower SE of estimates than BLUP SE estimates (Table 3, Additional Table 1). Hence, when not specified, results are given for GBLUP model in the rest of the paper and GBLUP results were mainly discussed.

**Table 3.** Genetic parameters estimated under GBLUP model for rainbow trout between B1 (juveniles) and B2 (harvest size).

|  | TLE | BW1 | BW2 | FL | Fat | HGC% |
| --- | --- | --- | --- | --- | --- | --- |
| TLE | **0.29 ± 0.05** | -0.49 ± 0.13 | 0.00 ± 0.12 | -0.02 ± 0.12 | 0.13 ± 0.10 | 0.11 ± 0.10 |
| BW1 | -0.07 ± 0.03 | **0.19 ± 0.04** | 0.45 ± 0.13 | 0.35 ± 0.13 | -0.05 ± 0.12 | -0.20 ± 0.11 |
| BW2 | NA | NA | **0.26 ± 0.04** | 0.75 ± 0.05 | -0.15 ± 0.10 | 0.08 ± 0.10 |
| FL | NA | NA | 0.86 ± 0.01 | **0.31 ± 0.04** | -0.25 ± 0.09 | 0.02 ± 0.09 |
| Fat | NA | NA | 0.10 ± 0.03 | 0.01 ± 0.03 | **0.45 ± 0.04** | 0.28 ± 0.08 |
| HGC% | NA | NA | 0.13 ± 0.03 | 0.14 ± 0.03 | 0.19 ± 0.03 | **0.61 ± 0.04** |

Heritability estimates in bold on the diagonal, genetic correlations on the upper triangle, phenotypic correlations on the lower triangle for standardised time to loss of equilibrium in acute hyperthermia challenge (TLE); BW1, body weight of batch 1; BW2, body weight of batch 2; FL, fork length; Fat%, fillet fat percentage; HGC% headed gutted carcass yield. All values are given ± standard error. NA: not applicable as traits were measured on different individuals.

BLUP and GBLUP models were tested with and without random dam effect for all traits. Random dam effect improved AIC of BLUP and GBLUP models for TLE but not for the other traits (data not shown) and was therefore only kept in TLE models.

Maternal effect was found to explain 7.0 ± 3.0 % of the TLE phenotypic variance in BLUP model and 6.0 ± 3.0 % in GBLUP.

Pedigree heritability and genomic heritability of TLE were estimated at similar values of 0.24 ± 0.07 and 0.29 ± 0.05, respectively. The two extreme groups in mean and SD, G1 and G7, had limited impact on the heritability of TLE. When ignoring G1 and G7 data, heritability estimates based on the remaining groups were 0.31 ± 0.05 and 0.28 ± 0.05, respectively. For this reason, G1 and G7 were kept in the dataset.

Heritability estimates were low for BW1 (0.19 ± 0.04), medium for BW2 (0.26 ± 0.04) and FL (0.31 ± 0.04) and high for Fat% (0.45 ± 0.04) and HGC% (0.61 ± 0.04).

As shown in Table 3, genetic correlation estimated between TLE and BW1 was clearly negative (-0.49 ± 0.13). In contrast, those estimated between TLE and BW2 and other production traits (FL, Fat% and HGC%) were all close to zero (between -0.02 ± 0.12 and 0.13 ± 0.10).

Phenotypic correlations were only estimated between traits measured within the same batch as it was impossible to estimate them between traits measured on different individuals. Phenotypic correlations are shown in the lower triangle of Table 3. Phenotypic correlation between TLE and BW1 was negative but close to zero: -0.07 ± 0.03.

### Genome-wide association study for acute hyperthermia resistance in juveniles

GWAS was performed for TLE using HD genotypes (∼420K SNP). The Manhattan plot of the QTL detection is shown in Figure 4. Seven QTL were detected with a logBF ≥6 for a peak SNP and located on five chromosomes. Among these seven QTL, the one detected on chromosome 30 was discarded as no other SNP had a logBF ≥3 in a sliding window of 200 kb around the peak SNP (Affx-1237752048). A visual inspection of the genotyping clusters for this peak SNP indicated that the genotyping of this SNP was of poor quality and that the high logBF value should be considered an artefact. Therefore, we only considered the six QTL named TLE2-1, TLE13-1, TLE13-2, TLE13-3, TLE14-1 and TLE17-1 (Table 4).

**Figure 4.**
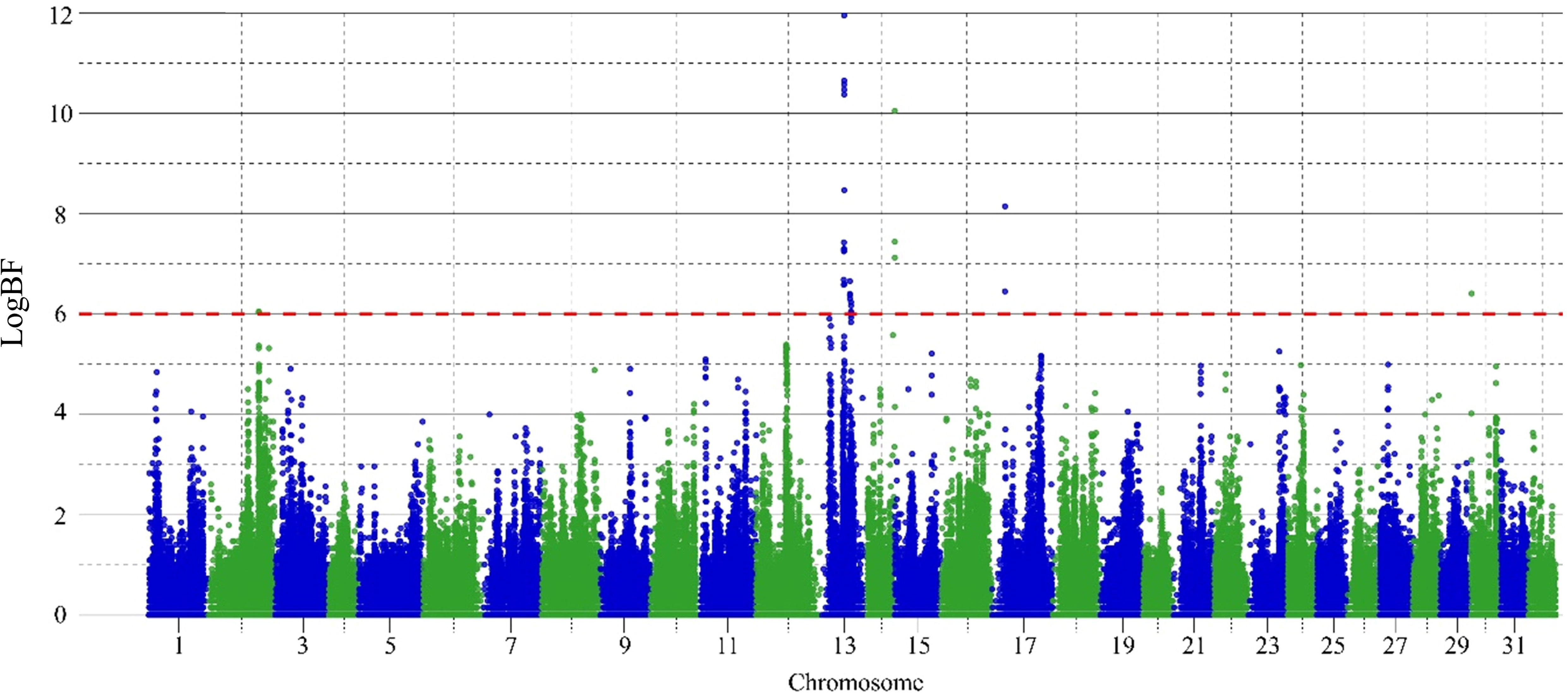
Manhattan plot of QTL detected for acute hyperthermia resistance trait in rainbow trout in Batch 1 (B1, juveniles). The red horizontal line corresponds to the QTL evidence threshold (logBF ≥6).

**Table 4.**
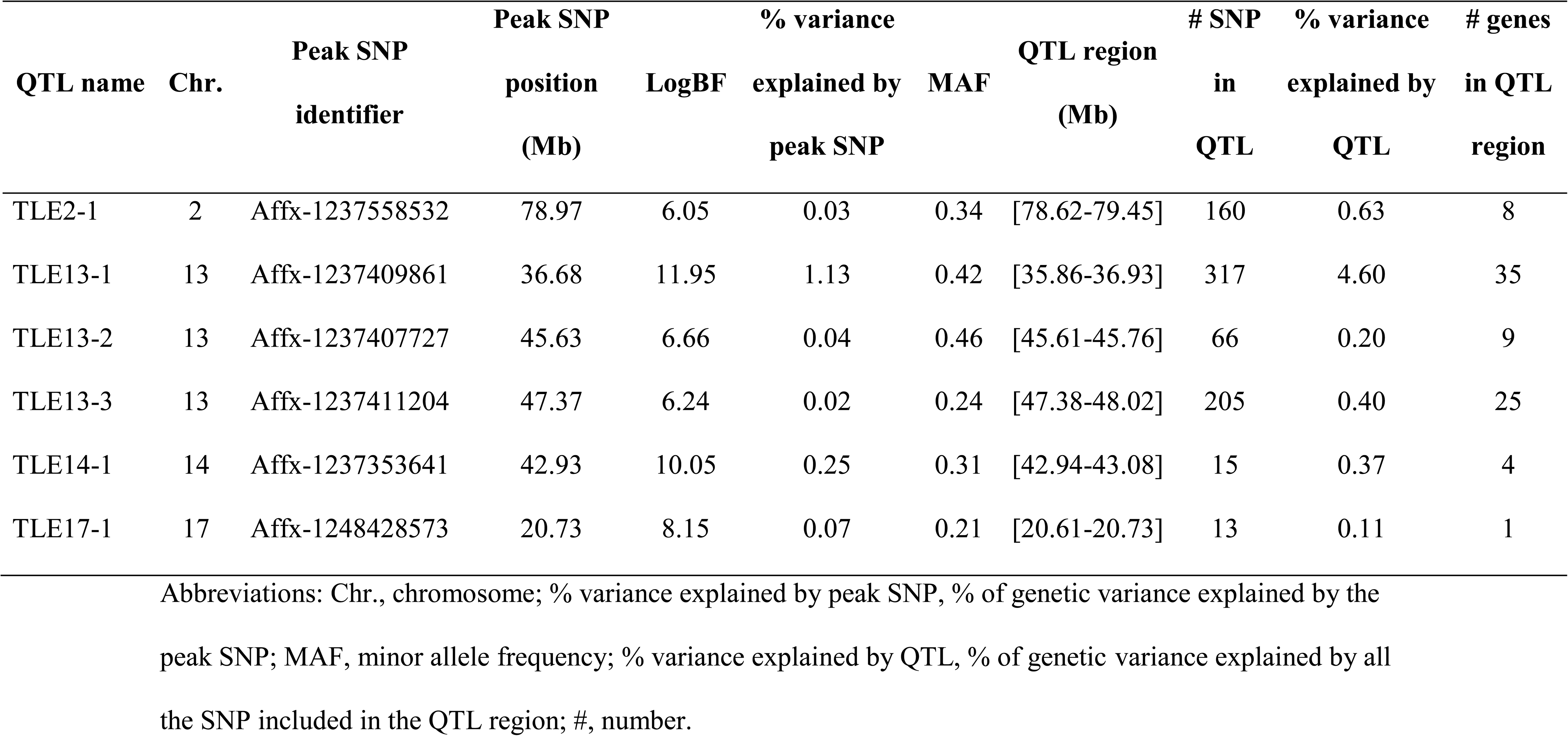
QTL effects and locations for acute hyperthermia resistance.

The peak SNP for TLE13-1 showed strong evidence for QTL with a logBF over 10. The whole QTL TLE13-1 explained more than 4% of the total genetic variance of TLE (Table 4). The five other QTL explained less than 1% of the genetic variance each (between 0.11% and 0.63%). This result suggests that resistance to acute hyperthermia is a highly polygenic trait.

Sizes of credibility intervals of QTL ranged between 0.12 and 1.07 Mb wide (Table 4). A total of 89 distinct genes were identified across all the QTL regions. The number of genes annotated in each QTL region is given in Table 4 and the gene names are reported in Additional table 2. Literature searches were carried out on genes in all QTL regions to identify genes that were previously associated with acute hyperthermia resistance. Meaningful candidate genes linked to acute hyperthermia resistance were found on QTL 13-1, 13-2 and 13-3 and will be discussed further.

### Comparison of TLE between genotypes at peak SNP

We analysed the individual TLE of fish, corrected for the dam and day effects, according to their genotypes at the peak SNP of each of the six identified QTL. Differences in TLE between the two homozygous genotypes were between 0.39 and 0.69 depending on SNP (Table 5), representing between 39% and 69% of the TLE phenotypic variation, as TLE was standardised to unity. As expected, the SNP for which the difference between the two homozygous genotypes was the strongest was for TLE13-1, the QTL with the highest logBF. Boxplots of TLE corrected for day and dam effects depending on the genotypes at the peak SNP of the 6 QTL are presented in additional figure 1.

**Table 5.**
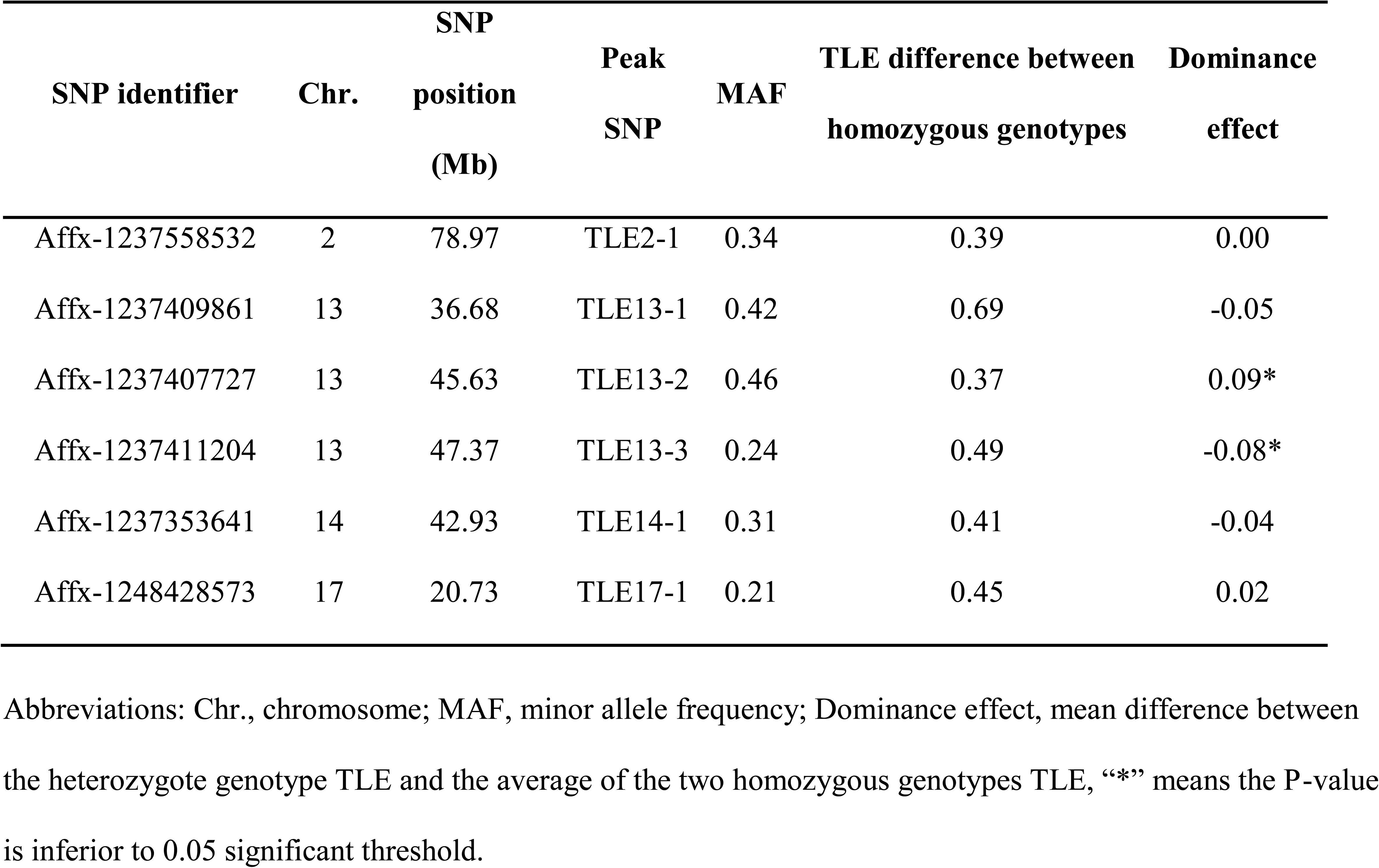
Difference in standardised acute hyperthermia resistance (TLE) for genotypes and dominance at the peak SNP.

| SNP identifier | Chr. | SNP<br>position<br>(Mb) | Peak<br>SNP | MAF | TLE difference between<br>homozygous genotypes | Dominance<br>effect |
| --- | --- | --- | --- | --- | --- | --- |
| Affx-1237558532 | 2 | 78.97 | TLE2-1 | 0.34 | 0.39 | 0.00 |
| Affx-1237409861 | 13 | 36.68 | TLE13-1 | 0.42 | 0.69 | -0.05 |
| Affx-1237407727 | 13 | 45.63 | TLE13-2 | 0.46 | 0.37 | 0.09* |
| Affx-1237411204 | 13 | 47.37 | TLE13-3 | 0.24 | 0.49 | -0.08* |
| Affx-1237353641 | 14 | 42.93 | TLE14-1 | 0.31 | 0.41 | -0.04 |
| Affx-1248428573 | 17 | 20.73 | TLE17-1 | 0.21 | 0.45 | 0.02 |

Abbreviations: Chr., chromosome; MAF, minor allele frequency; Dominance effect, mean difference between the heterozygote genotype TLE and the average of the two homozygous genotypes TLE, “\*” means the P-value is inferior to 0.05 significant threshold.

Significant dominance effects were observed for genotypes at peak SNP of TLE13-2 (p-value = 0.01) and TLE13-3 (p-value = 0.04) (Table 5).

### Refinement of QTL using isogenic lines data

Among the SNP present on the HD genotyping array, we identified 52, 68, 13, 45, 5 and 0 SNP respectively in TLE2-1, TLE13-1, -2, -3, TLE14-1 and TLE17-1 regions for which alleles differed between the sensitive and the resistance isogenic lines. Of all these SNP, three SNP located on TLE13-1 and TLE13-2 had a logBF ≥6 in the commercial population. Details about these SNP, their effects on the commercial population and the alleles of the resistant and sensitive lines are given in table 6. Consistently, the alleles held by the resistant isogenic line at these 3 SNP were predicted by the GWAS to have a favourable effect on the acute hyperthermia resistance in the selected population and vice-versa for the sensitive line.

**Table 6.**
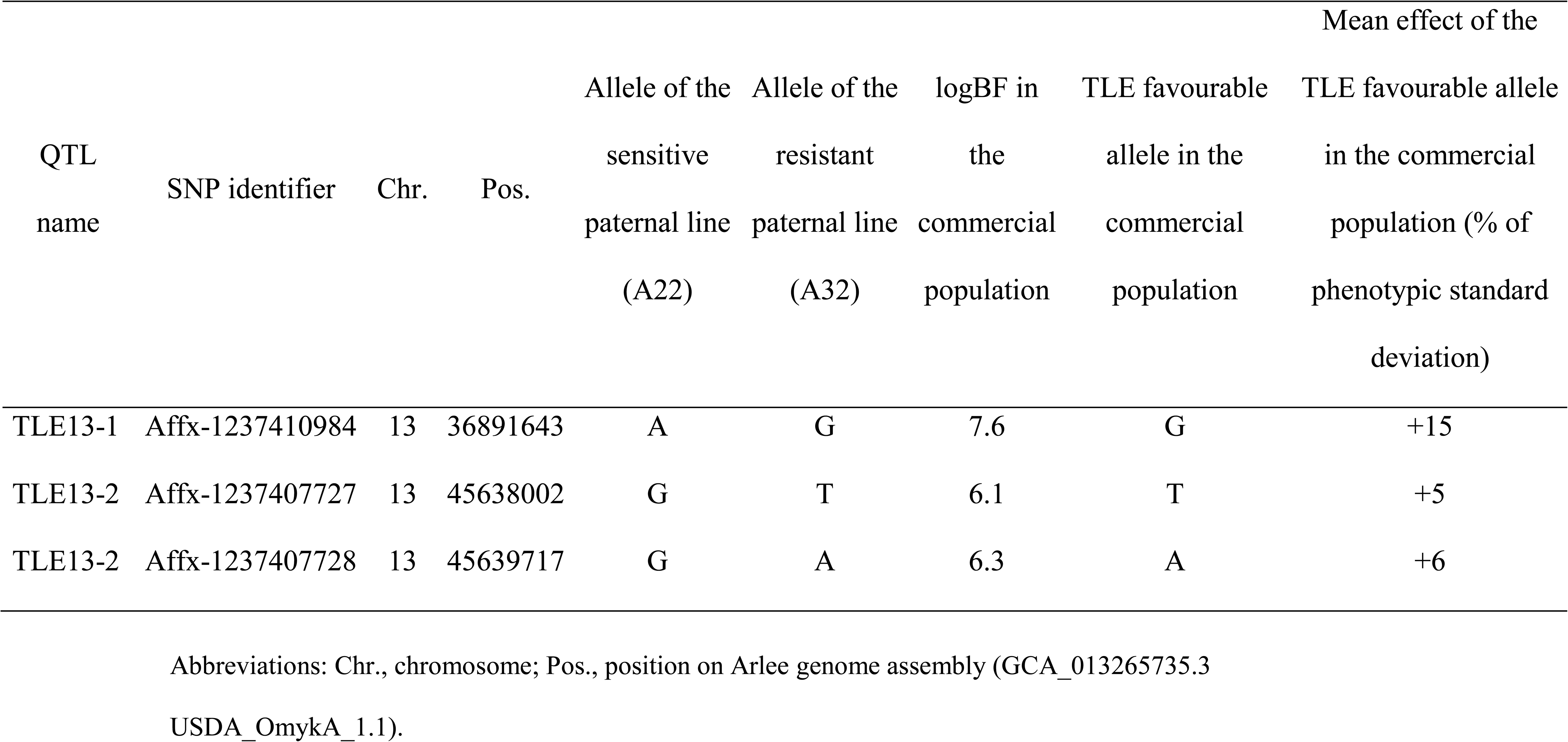
QTL shared between isogenic lines and commercial population.

## Discussion

In this study, we investigated the genetic architecture of resistance to acute hyperthermia in a French commercial population of rainbow trout at nine months of age. We estimated the genetic parameters of this trait and its correlations with production traits at a near 1 kg harvest weight (body weight, fork length, carcass yield, and fillet fat percentage). We also identified QTL for resistance to acute hyperthermia and searched for functional candidate genes.

### Defining acute hyperthermia resistance

In acute stress, the exposure to the stressor is short and intense while in chronic stress, the exposure is weaker and repeated over time [44]. However, this definition is context-dependent [45]. For example, in rainbow trout, acute hyperthermia resistance was defined as resistance to hyperthermia in a one-day challenge in Perry et al. [13], while it was in a one-week challenge in Chen et al. [46]. Whether acute or chronic, response to a stressor involves different physiological mechanisms [45,47,48]. Correlation between acute and chronic resistance to temperature was rarely studied but consistently, in rainbow trout and Atlantic salmon, no relationship was found between resistance to acute hyperthermia, measured as temperature at loss of equilibrium within a day, and resistance to chronic hyperthermia, measured as growth or survival at high temperature during more than a month [49, 50]. To our knowledge, no research was conducted on the relationships between acute resistance phenotypes to hyperthermia stress at different levels of exposure and intensity, for example between a one-day hyperthermia challenge and a two-day hyperthermia challenge. A better characterisation of the acute hyperthermia resistance phenotype, particularly its genetic and phenotypic correlations with semi-chronic and chronic hyperthermia resistances, would therefore be very valuable, as these stressful conditions might also happen in fish farms but is beyond the scope of this study. Because of this lack of information, in the rest of the discussion, we only considered hyperthermia stress to be acute if it made fish lose equilibrium within a day as we did and semi-chronic or chronic elsewhere.

### Phenotyping acute hyperthermia resistance

During acute hyperthermia resistance phenotyping challenges, water quality parameters were carefully checked to ensure they had limited impact on acute hyperthermia resistance of fish. O2 saturation was constantly maintained above 80% in acute hyperthermia challenges. At this level, it has been shown that O2 saturation does not interfere with resistance to acute hyperthermia in fish [51–53]. NH4+ concentration reached a maximum between 2 and 3 mg/L with a pH of 7.2 ± 0.4 six hours after the start of challenges. Given the pH and the short duration of the challenge, this NH4+ concentration remains under the toxicity threshold of rainbow trout [54]. CO2 concentration increased from 9 mg/L to 12 mg/L in the first three hours of the challenge. Chronic exposure for six months to a twice higher CO2 concentration (24 ± 1 mg/L) was reported to have no significant impact on rainbow trout growth and survival [55]. NH4+ and CO2 concentrations increases were therefore assumed to have minor effects on the resistance of fish to acute hyperthermia.

In the present paper, acute hyperthermia resistance was measured as the raw time before loss of equilibrium. In contrast, some other studies used cumulative degrees to quantify resistance to acute hyperthermia [13,16,56]. Cumulative degrees are calculated as a combination between time and temperature: the difference between temperature at each minute of the challenge and the initial temperature is cumulated from the beginning of the challenge to the time of loss of equilibrium [13]. Cumulative degrees phenotype is supposed to correct for differences in temperature increase rate between groups. However, in the present study, phenotypic correlation between cumulative degrees and raw time before loss of equilibrium was 0.98 and genetic correlation estimation did not converge as these two phenotypes were too correlated. This strong correlation might not be true in studies with more considerable variations in temperature arising between phenotyping groups compared to the present study. Because of this similarity between the two phenotypes, we decided to use the simplest one, time before loss of equilibrium.

There were significant disparities between phenotyping groups in terms of mean and SD of the rTLE. The number of acclimation days before challenge and the temperature six hours after the start of the challenge were strongly correlated with the groups rTLE means and SD and therefore may explain these disparities. However, these two factors being confounded, it was impossible to determine which of them had a predominant effect on acute hyperthermia means and SD of groups.

Acclimation, particularly the acclimation temperature which precedes acute hyperthermal stress, is a well-known factor influencing acute hyperthermia resistance of fish [10]. In the present study, first groups to be phenotyped were less acclimated to the river temperature (16- 18°C) which was higher than the temperature (11-12°C) in the fish farm before acute hyperthermia resistance phenotyping and during the transportation to the phenotyping platform (11-13°C). First groups to be phenotyped, thus less acclimated to 16-18°C higher temperature, had a lower mean in acute hyperthermia resistance. This was consistent with Chen et al. [57] which showed a strong positive correlation of 0.74 between acclimation temperature and acute hyperthermia resistance in rainbow trout.

The second identified factor was the temperature after six hours of challenge which was highly correlated with acute hyperthermia resistance means and SD of groups. Logically, phenotyping groups with a lower temperature after six hours of challenge resisted longer acute hyperthermia stress, as in Becker et al. [58]. Lower temperature increase rates could also increase acute hyperthermia resistance SD in ectotherms such as in the Cuyaba dwarf frog *Physalaemus nattereri* although this varies between species [59]. In the present study, we consistently found that phenotyping groups with a lower temperature after six hours of challenge had a higher mean and SD in acute hyperthermia resistance.

Group disparities were corrected by centring and reducing rTLE by group of challenge (TLE).

### Heritability estimates for acute hyperthermia resistance in juveniles and production traits

Pedigree- and genomic-based heritability estimates for acute hyperthermia resistance at 280 dpf were 0.24 ± 0.07 and 0.29 ± 0.05, respectively. These estimates are lower but consistent with an earlier study which reported a pedigree-based heritability of 0.41 ± 0.07 on a North-American population of commercial rainbow trout at an age between 148 and 300 dpf [13]. Our lower estimate of heritability is associated with the inclusion of the maternal effect, explaining 6-7% of TLE phenotypic variance. A maternal effect of similar magnitude was also detected for acute hypoxia resistance at 270 dpf in rainbow trout [60]. Ignoring the random maternal effect, the pedigree- and genomic-based estimates of heritability were increased to 0.39 ± 0.06 and 0.35 ± 0.04, i.e. similar to the estimate of Perry et al. [13]. We have no information on whether the significance of maternal effect was checked in Perry et al. [13].

The discovery of a significant maternal effect at age as late as 280 dpf was surprising as generally the magnitude of the maternal effect tends towards zero within the first year of life in fish for traits like growth or survival [61, 62]. It seems that the magnitude of maternal effects on acute hyperthermia resistance also shrinks with age in salmonids: maternal effects were found to explain 77% of acute hyperthermia resistance phenotypic variance in chinook salmon *Oncorhynchus tshawytscha* larvae weighing between 0.6 and 3.6 g [63] while it was not significant for acute hyperthermia resistance in Atlantic salmon at 297 dpf [64].

In chinook salmon larvae, there was a strong correlation between the average egg diameter of females and the average acute hyperthermia resistance of their offspring, suggesting that the significant maternal effect could be due to the size of the eggs [63]. In the present study, there was no sorting of eggs by size. It is therefore possible that the maternal effect comes from differences in mean egg size between females. Another hypothesis of the significant maternal effect could be the maternal intergenerational plasticity effects and more particularly their effects on mitochondria. Hence, acclimation of dams to high temperatures was shown to significantly affect the mitochondria respiration capacities of their offspring for up to at least 60 days in the sticklebacks *Gasterosteus aculeatus* [65]. Body weight of these offspring was shown to be influenced by the dam acclimation temperature, probably induced by the different mitochondria respiration capacities [65]. Still, in the sticklebacks, warmer temperatures experienced by dam were shown to influence mitochondrial DNA level in oocytes [66]. Mitochondrial DNA level in oocytes was shown to significantly affect growth and mortality at least in the first week of life [66]. It is therefore possible that maternal intergenerational plasticity could impact other traits such as acute hyperthermia resistance.

The heritability of BW1 was found in the low range of previous estimates for BW at similar ages (0.20-0.28) [67, 68]. Heritability estimates for all other production traits were also found consistent with those of a study on 17-month-old rainbow trout issued from another French commercial population of rainbow trout [69].

### Genetic correlations between acute hyperthermia resistance in juveniles and other traits

Genetic correlations were estimated between acute hyperthermia resistance and production traits commonly selected in fish breeding programs. The interest in estimating genetic correlations is twofold. First, from a biological point of view, the genetic correlations are likely to reveal the possible existence of shared biological pathways between traits [70]. Secondly, from a breeder’s point of view, the genetic correlations predict the influence of selecting one trait on responses for other traits of interest. Therefore, estimating genetic correlations is essential for describing the genetic architecture of a trait, as well as for optimisation of breeding programs.

The phenotypic correlation between TLE and BW1, both traits measured on B1 between 275 and 285 dpf, was close to zero but lightly negative (-0.07 ± 0.03). This result is consistent with the literature in which BW was reported to have a zero or negative effect on acute hyperthermia resistance in fish (reviewed in [71]).

The genetic correlation between TLE and BW1 was clearly negative (-0.49 ± 0.13). A previous study estimated a null genetic correlation (-0.03 ± 0.18) between acute hyperthermia resistance and body weight at 210-259 dpf in a North American population of rainbow trout [13]. This difference in result between the present study and the one from Perry et al. [13] might be due to genetic or environmental differences between the studied populations as was observed in Kause et al. for body weight [72], the different phenotyping ages or the method to produce the families by separated or mixed family rearing. However, our result is consistent with Debes et al. [64] which found in first-year Atlantic salmon a clearly negative genetic correlation of −0.86 ± 0.49 between acute hyperthermia resistance and fork length, fork length being highly correlated with body weight in Atlantic salmon [73] as well as in rainbow trout as shown in Table 3 and in Haffray et al. [32]. Trade-offs were suggested between body weight and acute hyperthermia resistance in rainbow trout [11,16,24] and in fish in general [71, 74], suggesting that there are physiological mechanisms underlying this negative relationship and thus potentially genetic basis. A controversial hypothesis to explain this link between body weight and acute hyperthermia resistance is the oxygen- and capacity-limitation of thermal tolerance. This theory states that the point of failure of acute hyperthermia resistance in fish is due to the inability of the organism to supply enough oxygen to the tissues as oxygen requirements increase exponentially with temperature [75]. Larger fish would be more sensitive to hyperthermia as their aerobic scope could be reduced compared to smaller fish [75]. However, this theory failed to predict acute hyperthermia resistance in some fish species and is therefore not enough by itself [51,76,77]. We also looked at whether QTL could be overlapping between TLE and BW1 and could explain some of this strong genetic correlation. Hence, TLE13-1, the most evident QTL associated with acute hyperthermia resistance was found in the same position as a QTL related to BW previously found in another population of rainbow trout at 410-481 dph [78]. In the present study, we found 1 QTL for BW1 and 12 QTL for BW2. Positions of BW QTL are given in Additional table 3. However, in our population, none of these QTL was at similar locations as the one identified in Ali et al. [78] or overlapped with any of the QTL associated with TLE we detected.

We estimated the genetic correlations between TLE and the production traits at near harvest age. The main selected trait in rainbow trout breeding programs is growth [15]. The genetic correlation between TLE and BW2 was not significantly different from zero. Selecting for acute hyperthermia resistance at nine months would therefore have no impact on growth at twenty months and vice versa.

Two opposing hypotheses may explain why genetic correlation is non-significant between TLE and BW2 while it is significant between TLE and BW1.

The first hypothesis would be that acute hyperthermia resistance is not a stable trait between nine and twenty months in rainbow trout. According to this hypothesis, fish resistance rankings to acute hyperthermia might change between ages and thus, genetic correlations estimated between production traits measured at twenty months and TLE measured at nine months might be poorly informative on the genetic correlations between production traits measured at twenty months and TLE measured at any other age than nine months. Genetic correlation between acute hyperthermia resistance at twenty months (not measured in the present study) and BW2 might still be strongly negative. Therefore, selecting for resistance to acute hyperthermia at nine months would not improve resistance throughout the life cycle of the fish, which would lose some of the interest in selecting for this trait. However, this is not the most likely hypothesis as several studies have shown good repeatability of acute hyperthermia resistance in brook trout Salvelinus fontinalis and rainbow trout over one year [16,56,79]. One study found no repeatability of acute hyperthermia resistance in sea bass *Dicentrarchus labrax* over eleven months [80]. However, it was in a semi-natural uncontrolled environment with strong genetic x environment interactions.

A second hypothesis would be that acute hyperthermia resistance is a stable trait with age and that the negative genetic correlation between acute hyperthermia resistance and body weight shrinks with age in rainbow trout. The stability of acute hyperthermia resistance trait with age is consistent with the studies cited in the previous paragraph. The shrinking genetic correlation between acute hyperthermia resistance and BW with age is consistent with Lagarde et al. [16] where body weight was found to have a significant effect on acute hyperthermia resistance in rainbow trout at nine months, but no effect at twenty months. If this assumption is correct, selecting for acute hyperthermia resistance at any age would have no impact on growth at twenty months and vice versa.

The genetic correlations between TLE and the other production traits (FL, Fat% and HGCW) were also close to zero. Therefore, selecting for TLE at nine months should have no impact on these traits and BW2 at twenty months. Also, current selection for production traits should not have impaired the acute hyperthermia resistance of rainbow trout.

### QTL associated with acute hyperthermia resistance in juveniles

In this study, we identified six QTL associated with acute hyperthermia resistance in a French commercial population of rainbow trout using a high-density chip with 665K SNP. QTL were found on chromosomes 2, 13, 14 and 17. Previous studies identified QTL associated with acute hyperthermia resistance on chromosomes 1, 9, 19 and sexual chromosome Y in a North-American population of rainbow trout using a limited number of markers composed of allozymes, RAPD and microsatellites [24–27,81]. The absence of common QTL between the present study and the previous ones is not surprising as population origins and marker densities are very different [82].

In total, the six QTL only explained 5% of the genetic variance of acute hyperthermia resistance (Table 4), of which the main QTL TLE13-1 explained 4% by itself. This suggests that resistance to acute hyperthermia is a highly polygenic trait in rainbow trout. Nevertheless, phenotypic differences according to genotypes at the peak SNP were substantial, with a mean TLE difference of up to 69% of phenotypic standard deviation between the favourable and unfavourable homozygotes. This phenotypic difference between homozygotes is considerable. In comparison, differences of 12% and 28% of phenotypic standard deviation between homozygotes at peak SNP for acute hyperthermia resistance were reported in the channel catfish and turbot *Scophthalmus maximus*, respectively [28, 83]. Thus, despite the low percentage of genetic variance explained at the population level, SNP peaks of the detected QTL seem to be good candidates for marker-assisted selection for acute hyperthermia resistance in the studied population.

In rainbow trout, previous studies have detected QTL associated with chronic hyperthermia resistance, i.e. the ability of fish to survive under chronic hyperthermia stress [46], and tolerance, i.e. the ability of fish to grow under chronic hyperthermia stress [84]. None of the QTL found in these two studies overlapped the QTL we detected in the present study. This result was expected as QTL tend to be population-specific, and also because resistance to acute hyperthermia and resistance or tolerance to chronic hyperthermia were shown to be distinct traits in salmonids [49, 50].

### Functional candidate genes implicated in acute hyperthermia resistance

In fish, GWAS for acute hyperthermia resistance were previously performed on the channel catfish [28] and the large yellow croaker [29]. These two studies identified 15 and 98 genes in the detected QTL regions respectively. Among these genes, one gene from the *dnaj* gene family (also called *hsp40* gene family) was systematically reported in the two studies: *dnajc25* in [28] and *dnajb4* in [29].

In line with these studies, we identified *dnajc7* at 150 kb from the SNP peak of the most significant QTL (TLE13-1). Dnaj proteins are co-chaperones of heat shock protein 70 (Hsp70), playing an important role in regulating this latter by recruiting Hsp70 partners and regulating the ATPase activity of the chaperone cycle [85]. In rainbow trout, *dnaj* family was found to be overexpressed in several organs (heart, brain, liver, spleen) during acute hyperthermia stress [86, 87] and individuals over-expressing *dnaj* genes were found more resistant to acute heat stress than fish with a lower level of expression [88, 89]. These results suggest that *dnaj* is a key gene family for resistance to acute hyperthermia in rainbow trout and other fish species.

Another member of *dnaj* family was also reported in a GWAS on chronic hyperthermia stress resistance in turbot with a time to loss of equilibrium higher than one week [83]. This result was surprising as there is growing evidence that acute and chronic hyperthermia resistances are distinct traits as previously mentioned in rainbow trout [49] and Atlantic salmon [50]. Nevertheless, it seems possible that some mechanisms of resistance to acute and chronic hyperthermia stress could be shared.

Still in TLE13-1 region, we identified other promising functional candidate genes. Indeed, very close to the peak SNP (less than 100 kb), we identified *hsp70b* and other *hsp70* homologues. *hsp70* genes family has several functions including molecular chaperoning and assisting the restoration or degradation of altered proteins [90]. *hsp70* genes are commonly associated with protein folding in generic response to stress exposure and more particularly acute hyperthermia exposure. In rainbow trout, larvae with a strong ability to upregulate the *hsp70b* gene were found significantly more resistant to acute hyperthermia compared to others [91] and in adults, *hsp70* was demonstrated to be upregulated during acute hyperthermia exposure [92, 93]. *hsp70b* was also shown to be the most over-expressed gene between a thermally selected strain and a thermal naïve strain of rainbow trout [89] and *hsp70b* was found to be the most over-expressed gene of the *hsp* family during acute hyperthermia conditions on immortalised rainbow trout gonadal fibroblasts [94]. *hsp70* was also demonstrated to have a role in acute hyperthermia resistance in other species. For example, in bay scallops *Argopecten irradians*, polymorphism in *hsp70* promoters significantly affected acute hyperthermia resistance, with *hsp70* being upregulated in more resistant individuals [95]. In the fruitfly *Drosophila buzzatii*, selection for heat resistance up to 64 generations has increased expression of *hsp70* [96].

*dnajc7* and *hsp70* are close genes, only 15 kb apart. Moreover, they were shown to interact in similar pathways and notably during acute hyperthermia stress in mammalians and fish [97–99]. These two properties suggest that these two genes may constitute a supergene associated with acute hyperthermia resistance, as previously identified for sex-specific migratory tendency in rainbow trout [100]. Supergenes are a group of segregated loci providing integrated control of complex adaptive phenotypes [101]. In other words, we hypothesise that *dnajc7* and *hsp70* have a role in acute hyperthermia resistance and that perhaps there are particularly favourable allelic combinations of these two genes.

We identified four other potential candidate genes for QTL TLE13-1: *nkiras2*, *cdk12*, *phb*, and *fkbp10*. These genes may be related to resistance to acute hyperthermia although less evident than *dnaj* and *hsp70*. The protein coded by *nkiras2* (NF-κB-inhibitor-interacting Ras 2) was shown to regulate NF-κB factors which are implied in homeostasis maintenance in mammalian cells [102] and in increasing the transduction of genes involved in inflammatory response in rainbow trout [103]. *nkiras2* was shown to be upregulated in gills of the chinook salmon exposed to acute hyperthermia stress (12°C to 25°C in three hours) [104]. cdk12 is a Pol II CTD kinase. In the fruitfly *Drosophila melanogaster*, cdk12 was shown to be involved in the control of the transcription of a set of genes involved in response to various stress factors: heat shock [105], DNA damage [106] and oxidative stress [107]. A lack of function of cdk12 was shown to increase the sensitivity of flies to oxidative stress [107]. Prohibitin (phb) is a highly conserved protein involved in many diverse functions such as chaperoning activities in mitochondria [108], the activation of transcription signalling pathways [109] or the regulation of cell survival and apoptosis [110]. *phb* was shown to be upregulated under acute heat stress in salt marsh mussel *Geukensia demissa* [111] and in a cell line derived from a human tongue squamous cell carcinoma [112]. These two studies disagree on the presumed role of prohibitin: Fields et al. (2016) hypothesised that prohibitin level increased in cells to delay or prevent heat-induced apoptosis while Jiang et al. (2013) argued that the increased abundance of prohibitin in cells may promote their apoptosis. Since prohibitin has many functions, some of which are poorly understood, it is difficult to know what effect this protein might have on resistance to acute hyperthermia in rainbow trout. *fkbp10* is part of the FK506-binding proteins genes family, involved in multiple functions including protein folding and repairing [113]. *fkbp10* was shown to be downregulated in salmonids subjected to chronic heat stress [114]. However, no study reported a differential expression of *fkbp10* during acute hyperthermia stress in fish.

In the isogenic lines, we found that the resistant line to acute hyperthermia was holding an allele which was also predicted to improve the resistance to acute hyperthermia in the commercial population in position 36.892 Mb of chromosome 13 while the sensitive isogenic line was carrying the deleterious allele. It seems that QTL TLE13-1 could be shared between the commercial population and isogenic lines, two distinct populations of rainbow trout with moderate genetic distance. Indeed, the fixation index (Fst) value was estimated at 0.09 [115] between the studied commercial population and the INRAE synthetic line from which isogenic lines were derived. Moreover, this new information, obtained from the whole-genome sequencing of the isogenic lines, refines the likely position of the TLE13-1 QTL candidate genes close to the position 36.892 Mb. This seems to confirm that genes near this position (*hsp70* genes family 36.744-36.827 Mb, *dnajc7* 36.842-36.851 Mb and *nkiras2* 36.851-36.853 Mb) are highly plausible functional candidate genes.

On TLE13-2, we identified two interesting genes previously found to be differentially expressed during acute hyperthermia exposure in fish: *ddx5* and *cygb1*. After acute hyperthermia stress, *ddx5* (probable ATP-dependent RNA helicase ddx5) was significantly downregulated in the salmonid taimen *Hucho taimen* [116] and *cygb1* (cytoglobin-1) was significantly upregulated in the channel catfish [117]. The resistant isogenic line was found to carry two alleles predicted to increase TLE in the commercial population. These two alleles are located in positions 45.638 and 45.640 Mb, which is closer to *ddx5* location (45.622-45.626 Mb) than *cygb1* one (45.711-45.717 Mb).

On TLE13-3, the gene *enpp7* (ectonucleotide pyrophosphatase/phosphodiesterase family member 7) was found close to the peak SNP. The protein from *enpp7* is an alkaline sphingomyelinase which hydrolyses membrane sphingomyelin to ceramide and phosphocholine [118]. We found no direct or indirect link between *enpp7* function and acute hyperthermia resistance but a GWAS of acute hyperthermia tolerance in pacific abalone *Haliotis discus* also found *enpp7* in a QTL region [119], indicating a possible role of this gene in acute hyperthermia resistance.

Last but not least, seven nuclear genes among the candidate genes were found to encode proteins with strong support of mitochondrial localisation in human according to the Human MitoCarta3.0 database (https://www.broadinstitute.org/mitocarta). These genes, some of which already discussed, were *pdhx* (TLE-2-1), *phb*, *fkbp10*, *acly*, *hsp70* (TLE-13-1), *elac2* and *sco1* (TLE-13-3). One limitation of acute hyperthermia resistance in fish may be related to the disrupted ability of mitochondria to produce ATP at high temperatures although evidence is still limited [120]. Interestingly, *pdhx* and *acly* genes are involved in ATP anabolic or catabolic processes. Pdhx, the pyruvate dehydrogenase protein X is part of the pyruvate dehydrogenase complex which catalyses the oxidative decarboxylation of pyruvate into acetyl-CoA, a reaction which notably ensures the link between glycolysis and the Krebs cycle [121, 122]. Acly, the ATP citrate lyase, is an enzyme involved in fatty acid biosynthesis, generating acetyl-CoA from citrate by consuming ATP [123, 124]. This also strengthens the hypothesis that the significant maternal effect found for TLE is related to mitochondria as mentioned above.

## Conclusion

This work provides new insights into the genetic architecture of acute hyperthermia resistance in rainbow trout juveniles using a novel high-density genotyping array with 665K markers. Heritability was moderate, and genetic correlations between acute hyperthermia resistance at juvenile stage and main production traits measured in sibs at harvest age were close to zero. Incorporating a trait for resistance to acute hyperthermia would make it possible to obtain robust animals without reducing production performance. However, this should be confirmed by ensuring that the genetic correlation between TLE at juvenile stage and harvest age is high. Our study demonstrated that acute hyperthermia resistance is polygenic in rainbow trout which is consistent with observations in other fish species. Indeed, the six identified QTL only explained around 5% of the genetic variance. The most significant QTL, located on chromosome 13, explained 4% of the genetic variance and genes directly associated with acute hyperthermia resistance (hsp70 genes family and dnajc7) were found close to the peak SNP of this QTL. QTL also contained genes associated with protein chaperoning, oxidative stress response, homeostasis maintenance and cell survival making them good candidates for further functional validation. As a preliminary validation of detected QTL, we investigated the genotypes of two isogenic lines showing contrasting resistance to acute hyperthermia. The resistant isogenic line was shown to carry favourable alleles in the region of two (including the main QTL) of the six QTL which shows that these QTL may be shared between distinct populations. Despite the polygenic architecture of acute hyperthermia resistance, the phenotypic mean difference between homozygotes at SNP peaks of QTL was strong, showing great potential for marker-assisted selection. All these results demonstrate the relevance of selective breeding to improve fish acute hyperthermia resistance.

## Declarations

### Ethics approval and consent to participate

The experiment was carried out according to the European guidelines; the protocols were evaluated and approved by the ethic committee ANSES/ENVA/UPC No 16 and authorised by the French ministry of higher education and research (APAFIS#24441-2020022417122193).

### Consent for publication

Not applicable

### Availability of data and materials

The data used for this research are not publicly available.

### Authors’ contributions

HL: Investigation, Resources, Methodology, Software, Formal analysis, Data curation, Validation, Visualization, Writing - original draft, Writing - review & editing. DL: Funding acquisition, Project administration, Supervision, Conceptualization, Writing - review & editing. PP: Project administration, Conceptualization, Investigation, Resources. MP: Software, Writing - review & editing. YF: Investigation, Resources. JA: Methodology, Software. ES: Investigation, Resources, Writing - review & editing. AAP: Investigation, Resources. FC: Project administration, Investigation, Resources. PH: Funding acquisition, Project administration, Writing - review & editing. AD: Resources, Methodology, Software. MDN Funding acquisition, Project administration Supervision, Conceptualization, Writing - review & editing. FP: Methodology, Software, Formal analysis, Data curation, Validation, Writing - review & editing.

## Acknowledgements

We are grateful to Yoann Cachelou from Viviers de Sarrance and Romain Morvezen from SYSAAF for their invaluable help in data collection.

## Additional files

**Additional file 1**

Format: word

Title: Boxplots of the centred and reduced acute hyperthermia resistance corrected from day and dam effects depending on the genotypes of the 1,328 fish at the peak SNPs of the six detected QTLs (see Table 5 for QTL characteristics).

Description: The Y-axis represents the TLE of fish (no unit) and the three colored boxes represent the three genotypes for a given SNP (two homozygous and one heterozygous). The dots represent the fish individual phenotype.

**Additional Table 1**

Format: Word

Title: Heritability and genetic correlations estimated under pedigree BLUP models.

Description: Heritability estimates in bold on the diagonal, genetic correlations on the upper triangle for resistance to acute hyperthermia measured as time to loss of equilibrium centred and reduced by group of challenge (TLE); body weight of batch 1 (BW1); body weight of batch 2 (BW2); fork length (FL); fillet fat percentage (Fat%); headed gutted carcass yield (HGC%). All values are given with their standard error.

**Additional Table 2**

Format: Word

Title: Candidate genes from the NCBI *Oncorhynchus mykiss* Annotation Release 100 (GCF_002163495.1) that were located within the QTL regions.

Description: none.

**Additional Table 3**

Format: Word

Title: QTL of body weight in B1 (BW1) and B2 (BW2).

Description: Abbreviations: Chr: chromosome; #, number; % variance explained by QTL, % of genetic variance explained by all the SNPs included in the QTL region.

