## Supplementary material for "Genetic architecture of acute hyperthermia resistance in juvenile rainbow trout (Oncorhynchus mykiss) and genetic correlations with production traits"

Additional figure 1. Boxplots of the centred and reduced acute hyperthermia resistance corrected from day and dam effects depending on the genotypes of the 1,328 fish at the peak SNPs of the six detected QTLs (see Table 5 for QTL characteristics). The Y-axis represents the TLE of fish (no unit) and the three colored boxes represent the three genotypes for a given SNP (two homozygous and one heterozygous). The dots represent the fish individual phenotype.

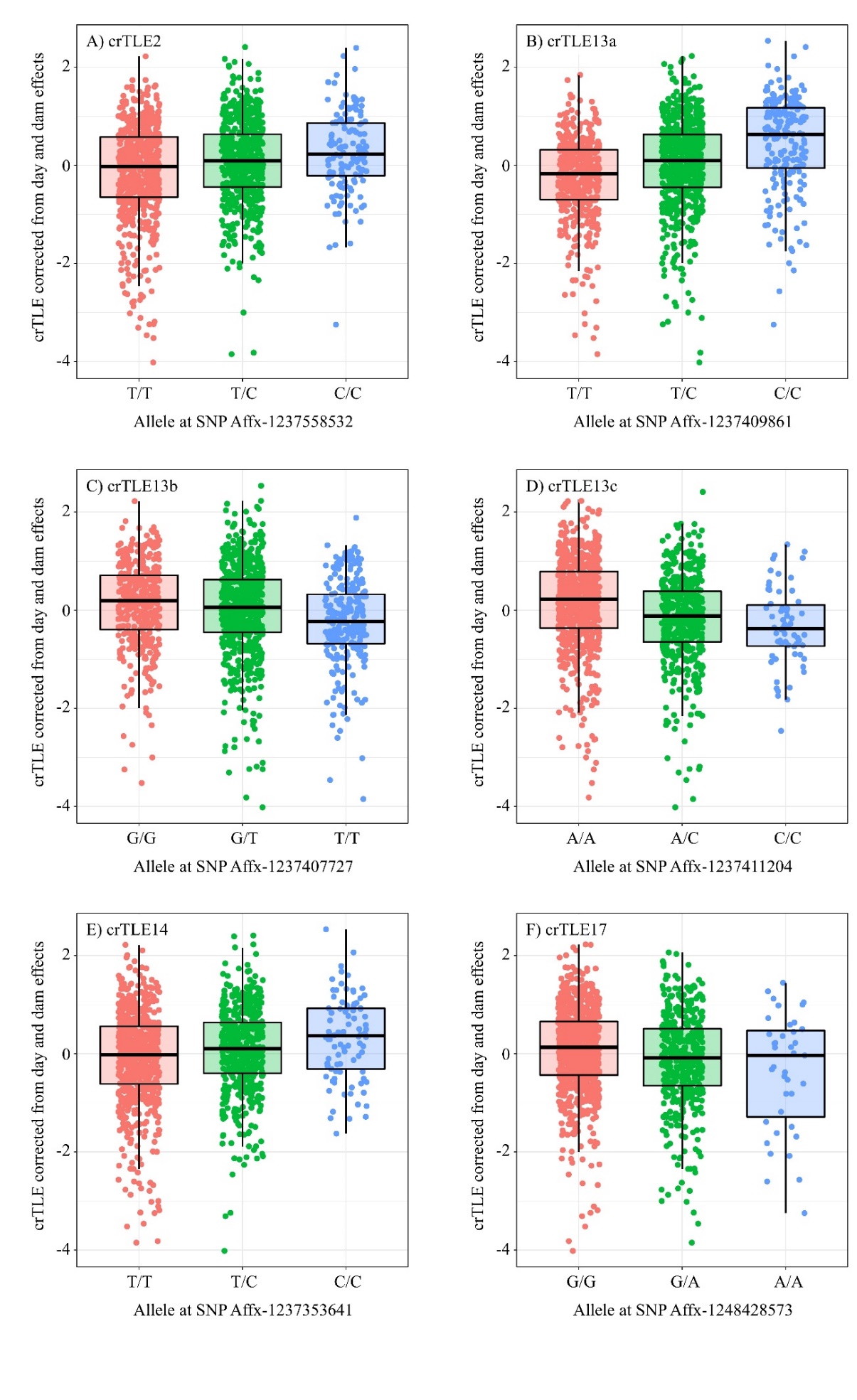

TLE17-1

TLE13-3

TLE13-1

TLE2-1

TLE13-2

TLE14-1

Additional table 1**:** Heritability and genetic correlations estimated under pedigree BLUP models

| Trait | TLE | BW1 | BW2 | FL | Fat | HGC% |
| --- | --- | --- | --- | --- | --- | --- |
| TLE | **0.24 ± 0.07** | -0.56 ± 0.20 | 0.09 ± 0.18 | 0.15 ± 0.17 | -0.02 ± 0.16 | 0.03 ± 0.16 |
| BW_C1 |  | **0.14 ± 0.04** | 0.61 ± 0.20 | 0.55 ± 0.19 | -0.13 ± 0.17 | -0.10 ± 0.17 |
| BW_C2 |  |  | **0.32 ± 0.06** | 0.79 ± 0.05 | -0.18 ± 0.12 | 0.04 ± 0.13 |
| FL |  |  |  | **0.34 ± 0.06** | -0.27 ± 0.12 | -0.08 ± 0.12 |
| Fat% |  |  |  |  | **0.59 ± 0.07** | 0.30 ± 0.10 |
| HGC% |  |  |  |  |  | **0.65 ± 0.07** |

Heritability estimates in bold on the diagonal, genetic correlations on the upper triangle for resistance to acute hyperthermia measured as time to loss of equilibrium centred and reduced by group of challenge (TLE); body weight of batch 1 (BW1); body weight of batch 2 (BW2); fork length (FL); fillet fat percentage (Fat%); headed gutted carcass yield (HGC%). All values are given with their standard error.

Additional table 2: Candidate genes from the NCBI *Oncorhynchus mykiss* Annotation Release 100 (GCF_002163495.1) that were located within the QTL regions.

| Chr. | QTL name | Start | Stop | GeneID | Locus | Protein Name | Gene symbol |
| --- | --- | --- | --- | --- | --- | --- | --- |
| 2 | crTLE-2-1 | 78713876 | 78853171 | 110497066 | LOC110497066 | low-density lipoprotein receptor class A domain-containing protein 3 isoform X2 | *ldlrad3* |
| 2 | crTLE-2-1 | 78927625 | 78995514 | 110497088 | LOC110497088 | tripartite motif-containing protein 44 isoform X2 | *trim44* |
| 2 | crTLE-2-1 | 79048412 | 79049695 | 110497116 | LOC110497116 | four-jointed box protein 1 | *fjx1* |
| 2 | crTLE-2-1 | 79103714 | 79150860 | 110497132 | LOC110497132 | inactive serine protease PAMR1 isoform X1 | *pamr1* |
| 2 | crTLE-2-1 | 79220052 | 79231353 | 110497158 | LOC110497158 | excitatory amino acid transporter 2 isoform X1 | *slc1a2* |
| 2 | crTLE-2-1 | 79220052 | 79231270 | 110497158 | LOC110497158 | excitatory amino acid transporter 2 isoform X2 | *slc1a2* |
| 2 | crTLE-2-1 | 79246003 | 79276658 | 110502122 | LOC110502122 | CD44 antigen | *cd44* |
| 2 | crTLE-2-1 | 79304459 | 79350048 | 110502133 | LOC110502133 | pyruvate dehydrogenase protein X component; mitochondrial-like | *pdhx* |
| 2 | crTLE-2-1 | 79387253 | 79390796 | 110497164 | *ehf* | ETS homologous factor isoform X1 | *ehf* |
| 13 | crTLE-13-1 | 35901477 | 35907852 | 100135854 | *st8sia6* | alpha-2,8-sialyltransferase 8F isoform X1 | *st8sia6* |
| 13 | crTLE-13-1 | 35903308 | 35907852 | 100135854 | *st8sia6* | alpha-2,8-sialyltransferase 8E isoform X4 | *st8sia6* |
| 13 | crTLE-13-1 | 35921395 | 35922846 | 110486220 | LOC110486220 | gap junction gamma-1 protein | *gjc1* |
| 13 | crTLE-13-1 | 35978998 | 35991940 | 110486221 | *eftud2* | 116 kDa U5 small nuclear ribonucleoprotein component | *eftud2* |
| 13 | crTLE-13-1 | 35998608 | 35999447 | 110485158 | LOC110485158 | probable phosphatase phospho1 isoform X1 | *phospho1* |
| 13 | crTLE-13-1 | 35998608 | 35999378 | 110485158 | LOC110485158 | probable phosphatase phospho1 isoform X2 | *phospho1* |
| 13 | crTLE-13-1 | 36010249 | 36024890 | 110486223 | LOC110486223 | zinc finger protein 652 | *znf652* |
| 13 | crTLE-13-1 | 36036597 | 36038508 | 110486224 | LOC110486224 | prohibitin | *phb* |
| 13 | crTLE-13-1 | 36042581 | 36047073 | 110486225 | LOC110486225 | uncharacterized protein LOC110486225 | *n/a* |
| 13 | crTLE-13-1 | 36053140 | 36089664 | 110486226 | LOC110486226 | synaptic vesicle membrane protein VAT-1 homolog | *vat1* |
| 13 | crTLE-13-1 | 36131832 | 36166410 | 110486228 | LOC110486228 | rho-related GTP-binding protein RhoN | *rnd2* |
| 13 | crTLE-13-1 | 36239763 | 36281846 | 110485159 | LOC110485159 | C-type mannose receptor 2 | *mrc2* |
| 13 | crTLE-13-1 | 36286050 | 36294981 | 110486229 | LOC110486229 | serine/threonine-protein kinase tousled-like 2 | *tlk2* |
| 13 | crTLE-13-1 | 36297254 | 36303119 | 110486230 | *mettl2a* | tRNA N(3)-methylcytidine methyltransferase METTL2 | *mettl2a* |
| 13 | crTLE-13-1 | 36308821 | 36331580 | 110486231 | LOC110486231 | integrin beta-3 | *itgb3* |
| 13 | crTLE-13-1 | 36338477 | 36338806 | 110486232 | LOC110486232 | reprimo-like protein | *rprml* |
| 13 | crTLE-13-1 | 36364332 | 36367430 | 110486233 | LOC110486233 | proline-rich protein 29 | *prr29* |
| 13 | crTLE-13-1 | 36375415 | 36376296 | 110486234 | LOC110486234 | telethonin | *tcap* |
| 13 | crTLE-13-1 | 36390573 | 36396558 | 110486235 | LOC110486235 | melanoregulin | *mreg* |
| 13 | crTLE-13-1 | 36400392 | 36407324 | 110486236 | LOC110486236 | stAR-related lipid transfer protein 3 isoform X2 | *stard3* |
| 13 | crTLE-13-1 | 36414833 | 36442644 | 110486237 | LOC110486237 | protein phosphatase 1 regulatory subunit 1B | *ppp1r1b* |
| 13 | crTLE-13-1 | 36459717 | 36460793 | 118938276 | LOC118938276 | neurogenic differentiation factor 2-like | *neurod2* |
| 13 | crTLE-13-1 | 36507143 | 36521970 | 110486239 | LOC110486239 | cyclin-dependent kinase 12 | *cdk12* |
| 13 | crTLE-13-1 | 36525971 | 36543609 | 110486240 | LOC110486240 | F-box/LRR-repeat protein 20 | *fbxl20* |
| 13 | crTLE-13-1 | 36561359 | 36577152 | 110486241 | LOC110486241 | SH3 and cysteine-rich domain-containing protein 2 | *stac2* |
| 13 | crTLE-13-1 | 36632896 | 36661946 | 110486244 | LOC110486244 | disintegrin and metalloproteinase domain-containing protein 11 | *adam11* |
| 13 | crTLE-13-1 | 36665852 | 36671261 | 110486245 | LOC110486245 | endoplasmic reticulum protein SC65 | *p3h4* |
| 13 | crTLE-13-1 | 36672452 | 36677094 | 110486247 | LOC110486247 | peptidyl-prolyl cis-trans isomerase FKBP10 | *fkbp10* |
| 13 | crTLE-13-1 | 36681083 | 36684455 | 110486248 | LOC110486248 | cytosolic 5\\'-nucleotidase 3 | *nt5c3a* |
| 13 | crTLE-13-1 | 36685135 | 36687456 | 110487537 | LOC110487537 | kelch-like protein 10 | *klhl10* |
| 13 | crTLE-13-1 | 36690152 | 36692504 | 110486249 | LOC110486249 | kelch-like protein 11 | *klhl11* |
| 13 | crTLE-13-1 | 36694322 | 36717836 | 110486250 | LOC110486250 | ATP-citrate synthase isoform X2 | *acly* |
| 13 | crTLE-13-1 | 36727743 | 36730526 | 110486251 | LOC110486251 | 2\\',3\\'-cyclic-nucleotide 3\\'-phosphodiesterase | *cnp* |
| 13 | crTLE-13-1 | 36735963 | 36736740 | 110486258 | LOC110486258 | tetratricopeptide repeat protein 25-like | *ttc25* |
| 13 | crTLE-13-1 | 36743455 | 36745389 | 110512845 | LOC110512845 | heat shock 70 kDa protein | *hsp70* |
| 13 | crTLE-13-1 | 36750387 | 36752321 | 118938280 | LOC118938280 | heat shock 70 kDa protein | *hsp70* |
| 13 | crTLE-13-1 | 36757270 | 36759204 | 118938281 | LOC118938281 | heat shock 70 kDa protein | *hsp70* |
| 13 | crTLE-13-1 | 36771036 | 36772970 | 118938282 | LOC118938282 | heat shock 70 kDa protein-like | *hsp70* |
| 13 | crTLE-13-1 | 36778096 | 36780030 | 118938279 | LOC118938279 | heat shock 70 kDa protein | *hsp70* |
| 13 | crTLE-13-1 | 36784977 | 36786911 | 100137015 | *hsp70b* | heat shock protein 70b | *hsp70b* |
| 13 | crTLE-13-1 | 36798965 | 36800899 | 118938283 | LOC118938283 | heat shock 70 kDa protein-like | *hsp70* |
| 13 | crTLE-13-1 | 36818530 | 36820464 | 118936524 | LOC118936524 | heat shock 70 kDa protein-like | *hsp70* |
| 13 | crTLE-13-1 | 36825392 | 36827326 | 110485160 | LOC110485160 | heat shock 70 kDa protein | *hsp70* |
| 13 | crTLE-13-1 | 36842225 | 36851292 | 110486253 | LOC110486253 | dnaJ homolog subfamily C member 7 isoform X1 | *dnajc7* |
| 13 | crTLE-13-1 | 36850669 | 36853454 | 110486255 | LOC110486255 | NF-kappa-B inhibitor-interacting Ras-like protein 2 isoform X1 | *nkiras2* |
| 13 | crTLE-13-1 | 36850679 | 36853454 | 110486255 | LOC110486255 | NF-kappa-B inhibitor-interacting Ras-like protein 2 isoform X2 | *nkiras2* |
| 13 | crTLE-13-1 | 36852482 | 36853454 | 110486255 | LOC110486255 | NF-kappa-B inhibitor-interacting Ras-like protein 2 isoform X3 | *nkiras2* |
| 13 | crTLE-13-2 | 45560259 | 45612271 | 110486546 | LOC110486546 | E3 ubiquitin-protein ligase SMURF2 | *smurf2* |
| 13 | crTLE-13-2 | 45621652 | 45626669 | 110486548 | LOC110486548 | probable ATP-dependent RNA helicase DDX5 isoform X1 | *ddx5* |
| 13 | crTLE-13-2 | 45621652 | 45626287 | 110486548 | LOC110486548 | probable ATP-dependent RNA helicase DDX5 isoform X2 | *ddx5* |
| 13 | crTLE-13-2 | 45632813 | 45646510 | 110486549 | LOC110486549 | envoplakin | *evpl* |
| 13 | crTLE-13-2 | 45652624 | 45653872 | 100135782 | LOC100135782 | retinal arylalkylamine N-acetyltransferase | *n/a* |
| 13 | crTLE-13-2 | 45657939 | 45677701 | 110486550 | LOC110486550 | sphingosine kinase 1-like | *sphk1* |
| 13 | crTLE-13-2 | 45680937 | 45692868 | 110486551 | LOC110486551 | fas-binding factor 1 isoform X4 | *fbf1* |
| 13 | crTLE-13-2 | 45680937 | 45692307 | 110486551 | LOC110486551 | fas-binding factor 1 isoform X5 | *fbf1* |
| 13 | crTLE-13-2 | 45680937 | 45691568 | 110486551 | LOC110486551 | fas-binding factor 1 isoform X6 | *fbf1* |
| 13 | crTLE-13-2 | 45710936 | 45716457 | 110486554 | LOC110486554 | cytoglobin-1 | *cygb1* |
| 13 | crTLE-13-2 | 45730762 | 45740418 | 110486555 | LOC110486555 | phosphoribosyl pyrophosphate synthase-associated protein 1 isoform X1 | *prpsap1* |
| 13 | crTLE-13-2 | 45730762 | 45739693 | 110486555 | LOC110486555 | phosphoribosyl pyrophosphate synthase-associated protein 1 isoform X4 | *prpsap1* |
| 13 | crTLE-13-2 | 45746133 | 45746759 | 110486556 | LOC110486556 | ras-related protein Rap-2a | *rap2a* |
| 13 | crTLE-13-2 | 46941309 | 47041211 | 110485195 | LOC110485195 | RNA binding protein fox-1 homolog 3-like isoform X2 | *rbfox3* |
| 13 | crTLE-13-3 | 47395433 | 47400215 | 110486591 | LOC110486591 | ectonucleotide pyrophosphatase/phosphodiesterase family member 7 | *enpp7* |
| 13 | crTLE-13-3 | 47403950 | 47407911 | 110486592 | LOC110486592 | chromobox protein homolog 2 isoform X1 | *cbx2* |
| 13 | crTLE-13-3 | 47432794 | 47433462 | 110486593 | LOC110486593 | noggin-3-like | *nog3* |
| 13 | crTLE-13-3 | 47461417 | 47466583 | 110486594 | LOC110486594 | phosphatidylcholine transfer protein | *pctp* |
| 13 | crTLE-13-3 | 47469699 | 47470130 | 110485196 | *tmem100a* | transmembrane protein 100 | *tmem100* |
| 13 | crTLE-13-3 | 47482712 | 47504517 | 110486595 | LOC110486595 | monocyte to macrophage differentiation factor isoform X1 | *mmd* |
| 13 | crTLE-13-3 | 47488516 | 47504517 | 110486595 | LOC110486595 | monocyte to macrophage differentiation factor isoform X2 | *mmd* |
| 13 | crTLE-13-3 | 47492036 | 47504517 | 110486595 | LOC110486595 | monocyte to macrophage differentiation factor isoform X3 | *mmd* |
| 13 | crTLE-13-3 | 47511898 | 47522430 | 110486596 | LOC110486596 | hepatic leukemia factor | *hlf* |
| 13 | crTLE-13-3 | 47566915 | 47597083 | 110486597 | LOC110486597 | forkhead box protein K2 | *foxk2* |
| 13 | crTLE-13-3 | 47612912 | 47613316 | 110487550 | LOC110487550 | phosphoinositide-interacting protein | *pirt* |
| 13 | crTLE-13-3 | 47639788 | 47706368 | 110486600 | LOC110486600 | protein shisa-6 isoform X3 | *shisa6* |
| 13 | crTLE-13-3 | 47806788 | 47814889 | 110486601 | LOC110486601 | zinc phosphodiesterase ELAC protein 2 | *elac2* |
| 13 | crTLE-13-3 | 47818946 | 47830360 | 110486602 | LOC110486602 | dual specificity mitogen-activated protein kinase kinase 4 isoform X4 | *map2k4* |
| 13 | crTLE-13-3 | 47831328 | 47832595 | 110486605 | LOC110486605 | transmembrane protein 220 isoform X1 | *tmem220* |
| 13 | crTLE-13-3 | 47833460 | 47835282 | 110486603 | LOC110486603 | protein SCO1 homolog, mitochondrial | *sco1* |
| 13 | crTLE-13-3 | 47875547 | 47911501 | 110486607 | LOC110486607 | growth arrest-specific protein 7 isoform X1 | *gas7* |
| 13 | crTLE-13-3 | 47892249 | 47911501 | 110486607 | LOC110486607 | growth arrest-specific protein 7 isoform X4 | *gas7* |
| 13 | crTLE-13-3 | 47894202 | 47911501 | 110486607 | LOC110486607 | growth arrest-specific protein 7 isoform X3 | *gas7* |
| 13 | crTLE-13-3 | 47920503 | 47921621 | 110486610 | LOC110486610 | recoverin-like | *rcvrn* |
| 13 | crTLE-13-3 | 47927253 | 47932236 | 110486609 | LOC110486609 | germ cell-specific gene 1-like protein | *gsg1l* |
| 13 | crTLE-13-3 | 47933613 | 47934227 | 110486608 | LOC110486608 | heparan sulfate glucosamine 3-O-sulfotransferase 3B1-like | *hs3st3b1* |
| 13 | crTLE-13-3 | 47948696 | 47953944 | 110486613 | LOC110486613 | dehydrogenase/reductase SDR family member 7C-B isoform X2 | *dhrs7cb* |
| 13 | crTLE-13-3 | 47948696 | 47952377 | 110486613 | LOC110486613 | dehydrogenase/reductase SDR family member 7C-B isoform X1 | *dhrs7cb* |
| 13 | crTLE-13-3 | 47948696 | 47950644 | 110486613 | LOC110486613 | dehydrogenase/reductase SDR family member 7C-B isoform X3 | *dhrs7cb* |
| 13 | crTLE-13-3 | 47948696 | 47950307 | 110486613 | LOC110486613 | dehydrogenase/reductase SDR family member 7C-B isoform X4 | *dhrs7cb* |
| 13 | crTLE-13-3 | 47953208 | 47958915 | 110486612 | LOC110486612 | splicing factor U2AF 65 kDa subunit | *u2af2* |
| 13 | crTLE-13-3 | 47962136 | 47983861 | 110486615 | LOC110486615 | myosin phosphatase Rho-interacting protein isoform X4 | *mprip* |
| 13 | crTLE-13-3 | 47962136 | 47981868 | 110486615 | LOC110486615 | uncharacterized protein LOC110486615 isoform X1 | *n/a* |
| 13 | crTLE-13-3 | 47986112 | 47988937 | 110486616 | LOC110486616 | transcription factor 20 | *tcf20* |
| 13 | crTLE-13-3 | 48002720 | 48010337 | 110486618 | LOC110486618 | serine/threonine-protein kinase SBK1 | *sbk1* |
| 14 | crTLE-14-1 | 42873811 | 42922424 | 110514351 | LOC110514351 | highly divergent homeobox | *hdx* |
| 14 | crTLE-14-1 | 42948335 | 43016754 | 110534813 | *rps6kal* | ribosomal protein S6 kinase alpha-6 isoform X1 | *rps6ka6* |
| 14 | crTLE-14-1 | 42977886 | 43016754 | 110534813 | *rps6kal* | ribosomal protein S6 kinase alpha-6 isoform X5 | *rps6ka6* |
| 14 | crTLE-14-1 | 43054693 | 43061444 | 110533241 | *hmgn6* | high mobility group nucleosome binding domain 6 isoform X2 | *hmgn6* |
| 14 | crTLE-14-1 | 43056027 | 43057409 | 118938746 | LOC118938746 | putative GPI-anchored protein pfl2 | *pfl2* |
| 17 | crTLE-17-1 | 20580154 | 20731773 | 110485646 | LOC110485646 | nuclear factor 1 X-type-like isoform X11 | *nfix* |
| 17 | crTLE-17-1 | 20580154 | 20714831 | 110485646 | LOC110485646 | nuclear factor 1 X-type-like isoform X5 | *nfix* |
| 17 | crTLE-17-1 | 20580154 | 20681477 | 110485646 | LOC110485646 | nuclear factor 1 X-type-like isoform X9 | *nfix* |
| 17 | crTLE-17-1 | 20582622 | 20731773 | 110485646 | LOC110485646 | nuclear factor 1 X-type-like isoform X21 | *nfix* |
| 17 | crTLE-17-1 | 20584300 | 20731773 | 110485646 | LOC110485646 | nuclear factor 1 X-type-like isoform X4 | *nfix* |

Additional table 3: QTL of body weight in B1 (BW1) and B2 (BW2)

Abbreviations: Chr: chromosome; #, number; % variance explained by QTL, % of genetic variance explained by all the SNPs included in the QTL region.

| **Trait** | **Chr.** | **Peak SNP identifier** | **Peak SNP position (Mb)** | **logBF** | **MAF** | **QTL position (Mb)** | **# SNP in QTL** | **% variance explained by QTL** |
| --- | --- | --- | --- | --- | --- | --- | --- | --- |
| BW1 | 12 | Affx-1237391021 | 22.82 | 6.17 | 0.45 | [22.63-22.86] | 96 | 0.19 |
| BW2 | 8 | Affx-1237862049 | 76.45 | 6.00 | 0.30 | [75.99-76.45] | 91 | 0.19 |
| BW2 | 10 | Affx-1248372316 | 2.141 | 7.05 | 0.47 | [2.141-2.447] | 15 | 0.20 |
| BW2 | 10 | Affx-88951716 | 4.283 | 6.10 | 0.34 | [4.254-4.558] | 27 | 0.18 |
| BW2 | 10 | Affx-88928954 | 65.44 | 6.92 | 0.41 | [65.20-65.61] | 82 | 0.12 |
| BW2 | 11 | Affx-1237365647 | 24.77 | 6.19 | 0.29 | [24.77-24.95] | 49 | 0.04 |
| BW2 | 18 | Affx-277301666 | 12.47 | 6.06 | 0.39 | [12.29-12.47] | 24 | 0.04 |
| BW2 | 18 | Affx-1237515271 | 14.00 | 6.93 | 0.27 | [13.73-14.21] | 79 | 0.17 |
| BW2 | 18 | Affx-1237515881 | 14.51 | 6.74 | 0.22 | [14.50-14.53] | 3 | 0.05 |
| BW2 | 19 | Affx-1248512873 | 5.431 | 6.06 | 0.37 | [5.431-5.431] | 1 | 0.02 |
| BW2 | 22 | Affx-1237602579 | 48.15 | 9.48 | 0.49 | [47.96-48.23] | 20 | 0.42 |
| BW2 | 27 | Affx-1237666662 | 14.52 | 6.10 | 0.32 | [14.21-14.52] | 92 | 0.08 |
| BW2 | 28 | Affx-1237688703 | 41.54 | 9.63 | 0.42 | [41.43-41.54] | 6 | 0.23 |
